# Genomic and common garden approaches yield complementary results for quantifying environmental drivers of local adaptation in rubber rabbitbrush, a foundational Great Basin shrub

**DOI:** 10.1101/2021.10.14.464430

**Authors:** Trevor M. Faske, Alison C. Agneray, Joshua P. Jahner, Lana M. Sheta, Elizabeth A. Leger, Thomas L. Parchman

## Abstract

The spatial structure of genomic and phenotypic variation across populations reflects historical and demographic processes as well as evolution via natural selection. Characterizing such variation can provide an important perspective for understanding the evolutionary consequences of changing climate and for guiding ecological restoration. While evidence for local adaptation has been traditionally evaluated using phenotypic data, modern methods for generating and analyzing landscape genomic data can directly quantify local adaptation by associating allelic variation with environmental variation. Here, we analyze both genomic and phenotypic variation of rubber rabbitbrush (*Ericameria nauseosa*), a foundational shrub species of western North America. To quantify landscape genomic structure and provide perspective on patterns of local adaptation, we generated reduced representation sequencing data for 17 wild populations (222 individuals; 38,615 loci) spanning a range of environmental conditions. Population genetic analyses illustrated pronounced landscape genomic structure jointly shaped by geography and environment. Genetic-environment association (GEA) analyses using both redundancy analysis (RDA) and a machine-learning approach (Gradient Forest) indicated environmental variables (precipitation seasonality, slope, aspect, elevation, and annual precipitation) influenced spatial genomic structure, and were correlated with allele frequency shifts indicative of local adaptation at a consistent set of genomic regions. We compared our GEA based inference of local adaptation with phenotypic data collected by growing seeds from each population in a greenhouse common garden. Population differentiation in seed weight, emergence, and seedling traits was associated with environmental variables (e.g., precipitation seasonality) that were also implicated in GEA analyses, suggesting complementary conclusions about the drivers of local adaptation across different methods and data sources. Our results provide a baseline understanding of spatial genomic structure for *E. nauseosa* across the western Great Basin and illustrate the utility of GEA analyses for detecting the environmental causes and genetic signatures of local adaptation in a widely distributed plant species of restoration significance.

## Introduction

Changing climate, invasive species, and human land use activities have altered plant communities worldwide (Chen et al., 2011; Kelly & Goulden, 2008; Parmesan & Yohe, 2003; Thuiller et al., 2005). Attempts to restore and preserve native communities are important for maintaining ecosystem function and conserving evolutionary processes (Reusch et al., 2005; Whitham et al., 2003), and restoration efforts are increasing in response to global initiatives (Stange et al., 2021). Successful ecological restoration requires knowledge of spatial genetic structure and an understanding of how local adaptation varies across landscapes (Hufford & Mazer, 2003; Knapp & Rice, 1994; McKay et al., 2005). The preservation of locally adapted populations is a primary goal of conservation (Broadhurst et al., 2008; Knapp & Rice, 1994; Weeks et al., 2011), and restoration success is thought to be influenced by the degree to which seed sources are locally adapted to the environmental conditions of restoration sites (Hufford & Mazer, 2003; McKay et al., 2005). Accordingly, decades of research based on phenotypic variation, often using common gardens, has characterized the genetic basis of phenotypic variation and adaptation to local environments across plant populations (Baughman et al., 2019; Clausen et al., 1940; Hereford, 2009; Langlet, 1971; Leimu & Fischer, 2008; Linhart & Grant, 1996). This information can also be used for climate change-aware seedings, with “prestoration” efforts informed by the relationship between genetic variation, phenotypic traits, and environmental variation (Butterfield et al., 2017).

In conservation or restoration contexts, population genetic data have traditionally been employed to characterize genetic diversity and differentiation across the landscape (Broadhurst et al., 2008; Harrisson et al., 2014). Genetic differentiation among populations can reflect independent evolutionary histories across variable ecological contexts, and genetic diversity itself is often used as a proxy for evolutionary potential which can predict population viability or restoration outcomes (e.g., Reynolds et al., 2012; Wernberg et al., 2018). To date, most inferences regarding local adaptation for restoration have used phenotypic data from common gardens (Baughman et al., 2019; Kawecki & Ebert, 2004; Langlet, 1971), but more recent efforts have used DNA sequencing data for this purpose (e.g., Massatti & Knowles, 2020; Shryock et al., 2017, 2021). High throughput sequencing has improved the ability to characterize the fine-scale genetic structure of populations (e.g., Larroque et al., 2019; Novembre et al., 2008; Wang et al., 2013), to describe the genetic signatures of adaptation (Cao et al., 2011; Li et al., 2018; McKown et al., 2014), and to link environmental variation to evolutionary processes (Forester et al., 2016; Storfer et al., 2018). While whole genome resequencing can be costly when applied to large numbers of individuals, reduced representation approaches (e.g., GBS, RADseq; Andrews et al., 2016; Parchman et al., 2018) can rapidly and inexpensively assay tens of thousands of genetic variants in a large number of individuals without relying on prior genomic resources. These approaches have shown utility for describing fine-scale population structure and evolutionary history relevant to evolutionary potential in changing environments (Massatti, Doherty, et al., 2018; Shryock et al., 2021), and for characterizing the genetic context and consequences of restoration efforts (Bragg et al., 2020; Dittberner et al., 2019; Jahner et al., 2019; Williams et al., 2014). Reduced representation sequencing has also increasingly been used to analyze the genomic signature and environmental drivers of local adaptation in non-model organisms (Hendricks et al., 2018; Savolainen et al., 2013) or in the context of restoration (Flanagan et al., 2018; Harrisson et al., 2014; Hoffmann et al., 2015).

When describing the genetic basis of adaptation, analytical approaches can control for confounding processes underlying spatial genomic structure (such as isolation-by-distance; IBD) and describe the more restoration-relevant contribution of environmental variation to divergence among populations (isolation-by-environment; IBE; Bradburd et al., 2013; Lotterhos & Whitlock, 2015; Wang & Bradburd, 2014). Genetic-environment association (GEA) analyses leverage environmental and population genomic data to detect allele frequency shifts coordinated with environmental variation across space (i.e., adaptation) while testing for the influence of specific environmental variables (Forester et al., 2018; Günther & Coop, 2013; Rellstab et al., 2015). In contrast to uninformed genome scans, GEA analyses can be relatively robust to confounding processes such as IBD and are relatively effective at detecting weaker, multilocus signals of selection (Capblancq et al., 2018; Forester et al., 2016, 2018; Lotterhos & Whitlock, 2015). A variety of methodologies (e.g., mixed-models, ordination, hierarchical Bayesian modeling, machine learning) have been implemented across a range of taxa and data types (reviewed in Forester et al., 2018; Rellstab et al., 2015). Recent studies have used GEA to identify the genomic signal of local adaptation and its environmental drivers in the context of restoration or to predict population responses to climate change (Bay et al., 2018; Brauer et al., 2016; Fitzpatrick & Keller, 2015; Jia et al., 2020; Lu et al., 2019; Perrier et al., 2018; Supple & Shapiro, 2018). While few studies have utilized both phenotypic and GEA based methods in such contexts (but see Carvalho et al., 2020; Fitzpatrick et al., 2021; Jordan et al., 2020; Mahony et al., 2020; Steane et al., 2014), congruent results from independent approaches offer more comprehensive evidence for local adaptation (de Villemereuil et al., 2016; Rellstab et al., 2015; Sork et al., 2013). More generally, approaches combining phenotypic, genomic, and environmental data allow a baseline understanding of how local adaptation varies across the landscape and provide important context for restoration as managers consider how genetic material (i.e., seed sources) will respond to different environments (Breed et al., 2019; Carvalho et al., 2021; Flanagan et al., 2018).

Here, we apply genomic and phenotypic methods within the Great Basin Desert of North America, a topographically and ecologically distinct region covering five western states (∼540,000 km^2^) and characterized by pronounced environmental and biological diversity (Davies et al., 2011; West, 1983). Massive areas of perennial shrubland communities have been converted to exotic annual grasslands due to fire, drought, and the introduction of invasive plants (Bradley & Mustard, 2005), making this among the most threatened ecosystems in North America. A member of the Asteraceae, rubber rabbitbrush (*Ericameria nauseosa*) is a foundational shrub species of these communities. It is broadly distributed across western North America (Anderson, 1984; Toft, 1995), and exhibits exceptional phenotypic diversity (22 named varieties; Anderson, 1984; Nesom & Baird, 1993). *E. nauseosa* displays a wide range of phytochemical variation (Hegerhorst, Weber, & McArthur, 1987; Hegerhorst, Weber, McArthur, & Khan, 1987; Weber et al., 1985), serves an important role in recolonizing disturbed sites, and is a critical late-season flowering resource for herbivore and pollinator communities (Hansen, 1986; McArthur et al., 1979; Ogle et al., 2007; Rogers, 1979). Studies of *E. nauseosa* have reported remarkable herbivore (e.g., 39 gall-forming species at a single site; Fernandes et al., 2000) and pollinator diversity (e.g., visits from 60% of fall flying bees [167/278 species] across a 4-year study; Carril et al., 2018). Understanding how genetic variation is distributed across geographic and environmental space and how it relates to phenotypic variation is particularly crucial for foundational species, such as *E. nauseosa*, as the degree of local adaptation in such plants often has extended community and ecosystem-level consequences (Grady et al., 2011; Hughes et al., 2008).

Here, we analyze the geographic distribution of genetic variation across *E. nauseosa* populations in the western Great Basin, using both DNA sequencing and common garden-based phenotypic data. This species is an excellent candidate for such work, as there has been a history of interest in the habitat value of *E. nauseosa* (e.g., Weber et al., 1985), but to-date, it has not frequently been included in large-scale restoration projects that could obscure historic patterns of adaptation and gene flow. We used reduced representation sequencing to quantify spatial genomic structure and to analyze the association between genetic, geographic, and environmental variation. Specifically, we quantified the extent to which population genetic structure is associated with geographic and environmental variation and used two GEA approaches to detect the genetic signature of local adaptation and to identify the environmental variables underlying it. Additionally, we quantified phenotypic variation in seed weight, emergence, and seedling traits for the same populations using a greenhouse common garden. We tested for local adaptation by analyzing phenotype-environment associations and quantified the degree to which environmental variables were dually implicated across phenotype-environment and genetic-environment association analyses. Finally, we quantified the relative predictive power of geographic, environmental, and genetic data sampled in the field for explaining phenotypic variation in the common garden. Our results provide a population genomic perspective on local adaptation to environmental variation in the western Great Basin and illustrate how genomic and phenotypic approaches can yield complementary results for analyses of local adaptation capable of informing restoration.

## Methods and Materials

### Population sampling and common garden experiment

*Ericameria nauseosa* consists of two subspecies (*nauseosa* and *consimilis*) that additionally encompass 22 named varieties (Anderson, 1995; Nesom & Baird, 1993). Our study focused on habitats within the western Great Basin that are inhabited primarily by ssp*. nauseosa* var. *hololeuca*, with one collection of ssp*. nauseosa* var*. speciosa*. While genetic differentiation among subspecies is apparent, varietal classifications within subspecies are largely not reflective of genetic differentiation, especially within the var. *speciosa* complex (Authors., unpublished data). We bulk-collected seeds from 17 localities (Fig. 1A) in the western Great Basin during the Fall of 2017 from a minimum of 50 plants per location. After a survey of sagebrush steppe communities in the region with similar dominant vegetation, these sites were selected for sampling because they hosted multiple native species that could potentially serve as restoration seed sources. Plants face multiple challenges in these cold deserts, including low annual precipitation (215.2–388.4 mm for all sites included here), year-to-year variation in precipitation, strong precipitation and temperature seasonality (most precipitation arrives during the coldest months, followed by hot, dry summers), and soils that vary in depth, texture, and salinity (West, 1983). Most of the dominant plant species are perennial (West, 1983), and establishment from seed can be challenging; seedling survival through periods of summer drought is a limiting life history stage for many species, including *E*. *nauseosa* (Donovan et al., 1993).

**Figure 1:**
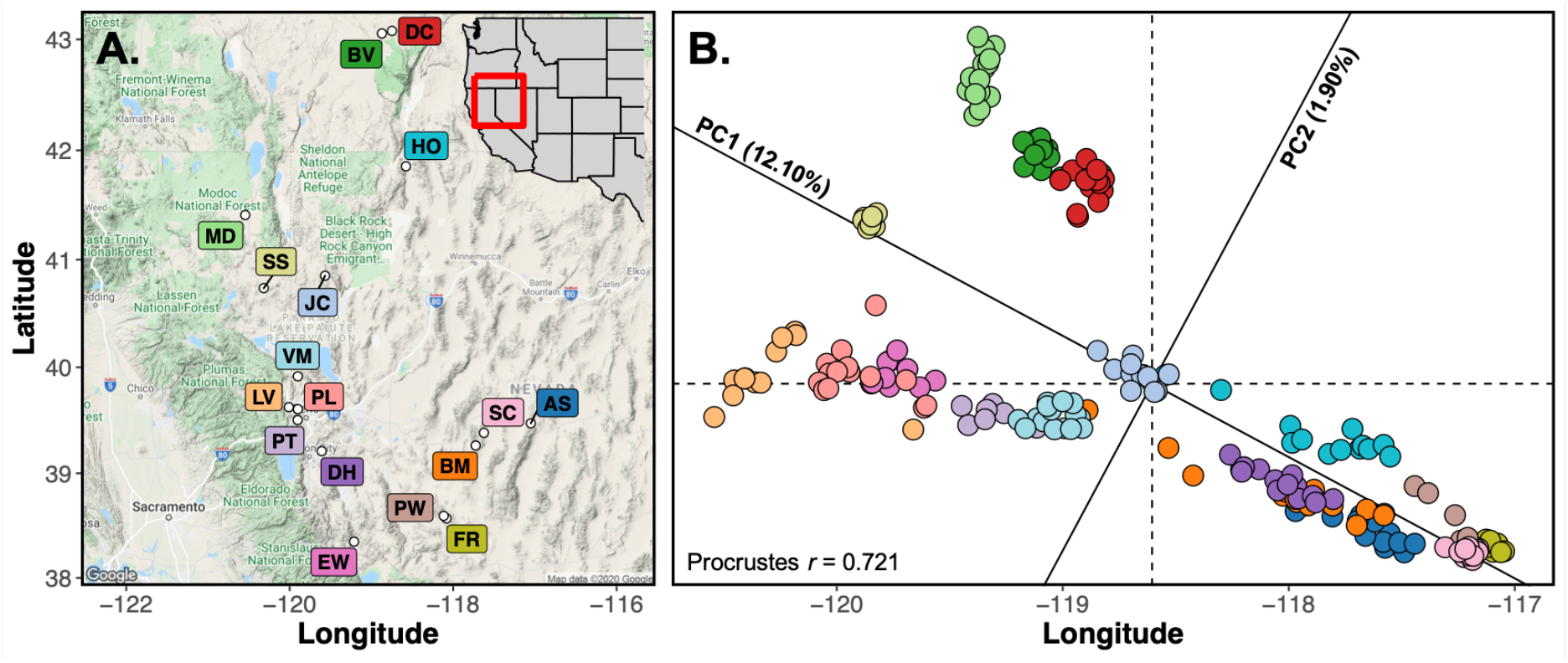
Landscape genomic structure strongly mirrors geography. (**A**) Map of the sampled localities for the 17 *E. nauseosa* populations. (**B**) Principal components analysis (PCA) of genotype probabilities illustrates spatial genomic structure. A Procrustes rotation of the first 2 PC axes onto latitude/longitude displays how spatial genetic structure is strongly predicted by geographic proximity. Colors of points in B match those of the population labels in A.

Adult leaf material was sampled from 15 plants per population and dried with silica gel to facilitate DNA extraction. Dried leaf material from each plant was ground into powder form using a Qiagen TissueLyser, and genomic DNA was extracted using Qiagen DNeasy Plant kits (Qiagen, Valencia CA). Extracted DNA was quality and quantity screened using a QIAxpert microfluidics electrophoresis device (Qiagen, Valencia CA).

We planted seeds from each sample locality in field soil in a greenhouse on the University of Nevada, Reno campus (latitude/longitude: 39.538, −119.805; elevation: 1370 m). Filled seeds (endosperm present) were separated from unfilled seeds, as they are visually distinct on a light table. We weighed ten batches of ten filled seeds per population to calculate an average seed weight per population. We then planted 100 filled seeds from each site following a randomized block design in the Fall of 2018. Seeds were monitored daily for emergence, and seedlings were harvested 40 days after emergence, using methods previously developed for describing seedling traits which are important predictors of success in restoration scenarios (e.g., Leger et al., 2020; Rowe & Leger, 2011). We separated seedlings into roots and shoots before drying and weighing. Samples were scanned and analyzed using WinRhizo imaging software (Regents Instruments, SianteFoy, QC, Canada). From these images we measured root length, root area, root and shoot mass, root mass ratio (root mass/total plant mass) and specific root length (length/mass). A descriptive summary of the full phenotypic dataset is available in Table S1. We generated genetic data from adult plants at each of the 17 sampling localities, while phenotypic data from the greenhouse common garden included only 14 of these localities where collected seeds were viable.

### Reduced representation sequencing and variant calling

Reduced representation libraries were prepared using a double-digest restriction-site associated DNA sequencing (ddRADseq) method (Parchman et al., 2012; Peterson et al., 2012). Genomic DNA was digested with two restriction endonucleases, *Eco*RI and *Mse*I, before uniquely barcoded Illumina adaptors were ligated to digested fragments using T4 DNA ligase. Following PCR amplification of pooled fragment libraries with a proofreading polymerase (Iproof; BioRad, Hercules CA), fragments ranging from 350 to 450 base pairs (bp) were size-selected using a Pippin Prep quantitative electrophoresis unit. Single-end sequencing (100 bp read lengths) was performed using two lanes on an Illumina HiSeq 2500 platform at the University of Wisconsin-Madison Biotechnology Center (Madison, WI).

After contaminant cleaning and demultiplexing, reads were assembled to a *de novo* reference with BOWTIE2 v 2.3.0 (Langmead & Salzberg, 2012), and variants were called using SAMTOOLS and BCFTOOLS v 1.8 (Li et al., 2009). After subsequent filtering with VCFTOOLS v 0.1.15 (Danecek et al., 2011), we retained bi-allelic single nucleotide polymorphisms (SNPs) with sequencing data for at least 70% of the samples and with minor allele frequency (MAF) > 0.05. To guard against genotyping bias stemming from the potential mis-assembly of paralogous regions, we removed loci with excessive coverage (summed depth across all samples *>* 10000), and those with abnormal heterozygosity (*F*_IS_ < −0.5 or *>* 0.5). Full methods for contaminant cleaning, assembly, variant calling, and additional filtering steps can be found in the Supporting Information.

### Population and landscape genomic variation

We used a hierarchical Bayesian model that incorporates genotype uncertainty (ENTROPY; Gompert et al., 2014; Shastry et al. 2021) to estimate genotype probabilities for each individual at each locus, infer the number of ancestral genetic clusters (*k*), and estimate individual ancestry coefficients (*q*). This model uses an allele frequency prior and incorporates genotype uncertainty arising from sequencing and alignment error as well as stochastic variation among individuals and loci in sequencing depth during parameter estimation (Gompert et al., 2014). We used genotype likelihoods calculated from SAMTOOLS as input, and used linear discriminant analysis, following *k*-means clustering of principal component scores to provide starting values of ancestry coefficients (*q*) to seed the MCMC (cf. Gompert et al., 2014). Genotype probability models with different ancestral clusters (*k*) from ENTROPY were estimated based on 100,000 MCMC iterations with a burn-in of 30,000 and thinning every tenth step. As the *k* = 2 model fit the data best (i.e., had the lowest deviance information criterion [DIC] value), genotype probabilities from this model were used for subsequent analyses (see Table S2 for DIC comparisons across *k* = 2–8). Due to complex fine-scale population structure, we do not present ancestry coefficient bar plots as they do not usefully illustrate population differentiation across the landscape. Rather, we used this model primarily to 1) obtain genotype probability estimates and thereby allow the incorporation of genotype uncertainty into subsequent analyses; and 2) generate individual ancestry coefficients (*q*), which were used to mitigate the influence of neutral population genetic structure in GEA analyses (see below).

We used principal component analysis (PCA), following standardization proposed by Patterson et al., (2006), to visualize population structure. A Procrustes rotation of the first two PC axes onto latitude and longitude was used to illustrate the concordance of genetic variation and geography (Wang et al., 2010), using the VEGAN R package (Oksanen et al., 2019; R Core Team, 2020). We characterized genetic differentiation among all populations using multilocus *F*_ST_ calculated with the HIERFSTAT R package (Goudet & Jombart, 2020) and population pairwise Nei’s *D* (Nei, 1972) calculated with custom R code. Isolation-by-Distance (IBD; Wright, 1943) and Isolation-by-Environment (IBE; Wang & Bradburd, 2014) were assessed using Mantel tests (Mantel, 1967) to associate genetic distance (Nei’s *D*) with geographic and environmental distances (see environmental description below). Geographic distances among populations were calculated as haversine distance using the GEOSPHERE R package (Hijmans, 2017). Using methods that incorporate genotype uncertainty implemented in ANGSD (Korneliussen et al., 2013, 2014), we estimated nucleotide diversity (*θ_π_*; Nei & Li, 1979) for each population and individual inbreeding coefficients (*F*) using NGSF (Vieira et al., 2013). See the Supporting Information for full description of ANGSD and NGSF methods.

In addition to the Mantel tests, IBD and IBE were estimated using a piecewise structural equation model (piecewiseSEM; Lefcheck, 2016; Shipley, 2000), as this allowed for estimation of the relative effect of environmental and geographic distance on genetic distance while accounting for covariance between the two (see Wang et al., 2013). The benefit of piecewiseSEM over other SEM approaches is it allows for local estimation of path coefficients to include random effects and varying sampling distributions. While many matrix correlation analyses use a permutation approach (i.e., Mantel or partial Mantel tests) to control for non-independence and test significance, *P*-value estimates from these approaches can be confounded by spatial autocorrelation among the matrices even when controlling for neutral processes (see Guillot & Rousset, 2013). Here we instead focus purely on relative effect sizes (or beta coefficient estimates). Population identification for each pairwise comparison was included in the model as a random effect to account for the correlated error structure inherent in pairwise observations, as suggested by Clarke (1993). All distance matrices were vectorized, centered, and standardized prior to analysis. Model fit was assessed with explained variance (marginal and conditional *r*^2^) and implemented using LME4 and PIECEWISESEM packages in R.

### Genetic-environment association analyses

We quantified environmental variation across sampling sites based on elevation and 30 climate variables calculated using the 30-year PRISM normals (1981-2010, 800m × 800m resolution) and the Climatic Water Deficit Toolbox (Dilts et al., 2015; Lutz et al., 2010). An R script provided by Redmond (2019) converts monthly normals to estimates of potential evapotranspiration, actual evapotranspiration, soil water balance, and climatic water deficit, which have been shown to effectively predict aspects of spatial and distributional variation across plant communities (Barga et al., 2018; Stephenson, 1998). Before any analyses, the pool of variables was reduced from 31 to ten after removing highly correlated variables (based on Pearson’s |*r*| ≤ 0.60) to control for multicollinearity. These ten variables represented variation in temperature, precipitation, elevation, evapotranspiration, and soil water capacity as well as precipitation seasonality, which is the degree of variability in monthly rainfall throughout the year (Walsh & Lawler, 1981; Table S3). Euclidean distances of the ten variables were used to represent environmental distance among populations where appropriate for distance-based analyses. Range values of the ten chosen environmental variables are presented in Table 2, and correlations of these variables with phenotypic measurements are illustrated in the Supporting Information (Figs. S1 and S2) while full summary statistics of all 31 variables and descriptions are provided in the Supporting Information (Table S3).

We quantified the degree to which genetic variation is associated with geographic and environmental variation using two approaches: (1) we estimated the proportion of genomic variance explained by environmental variation and the individual contributions of each environmental variable; and (2) we quantified locus-specific allele frequency shifts correlated with environmental variation to detect selection on genome variation and quantify the extent different variables underlie these shifts.

Two different GEA methods, redundancy analysis (RDA; Legendre & Legendre, 2012) and Gradient Forest (GF; Ellis et al., 2012; Fitzpatrick & Keller, 2015), were used to identify loci with allele frequency shifts potentially reflecting adaptation to environmental variation. Recent studies (Capblancq et al., 2018; Forester et al., 2016, 2018) have shown through both simulations and empirical data that RDA performs best (i.e., less prone to false negatives and false positives) in identifying loci under selection while remaining robust to complex demographic histories. We first used a partial RDA to assess the degree adaptive genetic variation among individuals is explained by a particular set of environmental variables, and to identify outlier loci potentially under selection. GF is a multivariate extension of the machine-learning algorithm, Random Forests (Breiman, 2001), originally developed to model communities of species assemblages. GF uses split functions along environmental gradients to predict both genome-wide and locus-specific allele frequency shifts (Fitzpatrick & Keller, 2015; Keller et al., 2018; reviewed by Fenderson et al., 2019). To reduce the confounding effects of neutral genetic variation on GEA inference, individual ancestry coefficients (*q*; generated by ENTROPY) were partialled out by regressing *q* from each environmental variable prior to analyses. This is statistically equivalent to conditioning within RDA (Legendre et al., 2011) but allowed partial effects to be accounted for in the same manner across both GEA analyses (RDA and GF).

To identify loci associated with environmental variation and potentially under selection, RDA was used with stringent outlier filtering of *±*3.5 SD on the first RDA axis (*P <* 0.0005; following Forester et al., 2018). These loci were then associated with individual environmental variables using Pearson’s correlation (*r*) to identify the strongest predictor. GF also gives a relative importance metric (*R*^2^) for each locus (i.e., loci with *R*^2^ > 0 indicate potential adaptive shifts in allele frequencies across a given environmental gradient) and the associated environmental variable. Results from RDA and GF analyses were compared using the correlation between the first RDA axis loading and GF *R*^2^ for environmental variables and outlier loci, and by quantifying the degree to which loci were associated with the same environmental predictor variables. To further show that these associations were not due to chance alone, a permutation test (*n* = 1000) was implemented in GF to verify there was a greater number of overlapping outlier loci between the two methodologies in the observed dataset than in random subsets of loci of equal size. Absolute values of the RDA loadings were used for any comparative analysis because only the relative strength of association was being assessed, not directionality. Most analyses and data manipulation were performed using the base STATS library or custom R functions, the VEGAN library (Oksanen et al., 2019) was used for the RDA, and the GRADIENTFOREST library (Ellis et al., 2012) was used for GF.

### Phenotypic variation

To complement GEA based inference of local adaptation and its environmental drivers, we measured phenotypes from field-collected seeds of 14 of the above populations in a greenhouse common garden, focusing on seed and seedling traits that have been shown to affect seedling establishment, an important component of restoration, in other taxa (Leger et al., 2020). Because seeds were collected from wild plants growing in different maternal environments, phenotypic variation likely represents both genetic and maternal environment effects, which is common in studies on local adaptation or provenance trials in wild plants (Baughman et al., 2019; Hereford, 2009; Risk et al., 2021). *Ericameria nauseosa* requires at least three years to produce seed, precluding a multigenerational design to control for maternal effects. The variables used for our analyses included a subset of traits with correlations of |*r*| ≤ 0.60 to control for multicollinearity. These included: seedling emergence (the number of seedlings that emerged and survived until harvest at 40 days; 92% of emerged seeds survived for 40 days, and thus for simplicity, we combined emergence and survival here), days to emergence, seed weight (average of ten seeds weighed in ten batches per population), shoot biomass, and average root diameter. Shoot biomass was chosen as the representative trait for seedling size as it is highly correlated (*r* = 0.62 – 0.92) with additional traits (total biomass, root length, root area, etc.) that are likely to be important in establishment (Caruso et al., 2019). Additionally, seedling emergence, seed weight, and shoot biomass were all highly positively correlated (*r* = 0.760 – 0.948) and were collapsed to a single composite variable using the first principal component axis, capturing 90.07% of the variance. This single composite variable was used for any multivariate analyses while the mean values for each trait in each population were used for univariate analyses.

The extent to which phenotypic variation might be influenced by genetic, environmental, and geographic variation was evaluated across a variety of analyses. First, we asked if traits measured in the common garden varied among populations using independent generalized linear mixed models. In these models, block was included as a random effect, a Bernoulli logistic regression was used for seedling emergence, a Poisson distribution was used for days to emergence, and Gaussian distributions were used for seed weight, shoot biomass, and average root diameter. Next, we analyzed the association of specific environmental variables with phenotypic variation from the common garden, asking whether there were phenotype-environment correlations consistent with local adaptation (Endler, 1986). For phenotype-environment associations, we used Random Forests (RF; Breiman, 2001) to assess potential relationships between phenotype and the same 10 environmental variables considered for the GEA analyses.

While phenotype-environment associations can be influenced by genetic drift, assessing how phenotypic variation relates to genetic variance can provide insight into the presence and mode of selection through *P*_ST_ - *F*_ST_ comparisons (McKay & Latta, 2002; Merilä & Crnokrak, 2001). *P*_ST_, an analogous proxy for *Q*_ST_ when pedigree information is unavailable (Brommer, 2011), was estimated for each phenotype and compared to multilocus *F*_ST._ We calculated *P*_ST_/*F*_ST_ across a range of *c/h*^2^ (where *c* is additive genetic variance, and *h^2^* is narrow sense heritability) to determine the critical value for *c/h*^2^ (i.e., where the lower 95% CI of *P*_ST_ equals the upper 95% CI of *F*_ST_) for each phenotype. More detailed methods and justifications for this analysis are available in the Supporting Information. To further evaluate congruence of environmental signal across genetic (GEA) and phenotypic (common garden) inference of local adaptation, we assessed whether specific environmental variables ranked similarly in their magnitude of association in both analyses. Importance (*R*^2^) from RF for the environmental variables on the mean trait values of seedling emergence, seed weight, days to emergence, shoot biomass, and average root diameter were correlated to the same environmental variable loading/importance from the GEA analyses (both RDA and GF).

Finally, the extent to which population-level phenotypic differences were associated with genetic, environmental, and geographic variation was assessed using a variance partitioning technique to estimate individual and shared contributions of each variable (Borcard et al., 1992). This approach uses partial RDAs to estimate the proportion of explained variance for each predictor variable, independently and combined, out of the total explained variance. Each model included phenotype as the response variable and all combinations of genetic, environmental, geographic variation as predictors. Phenotype was characterized by three variables: the composite variable of seedling emergence, seed weight, and shoot biomass, and the mean trait value of days to emergence and average root diameter for each population. Genetic and environmental data were collapsed using PCA and the first four PC axes were treated as composite variables to represent each group of data. Environmental data inputted into the PCA was the standardized raw data of all ten variables, while genetic data was represented by the variance-covariance matrix of the genotype probabilities, with the four PC axes explaining 86.0% and 62.7% of the total datasets, respectively. Geography was characterized by the latitude and longitude of the population. The total phenotypic variance explained (*PVE*) by the predictors and adjusted *r*^2^ (i.e., individual contribution in terms of total *PVE*) was estimated with the *varpart* function in the VEGAN package in R. Model significance, when appropriate, was assessed using independent RDA and pRDAs, permutation-based ANOVAs (*n* = 999), and significance thresholds of *α* ≤ 0.05. All measurements were centered and standardized prior to analyses.

## Results

### Population and landscape genomic variation

Two lanes of Illumina sequencing produced ∼613 million reads across 222 individuals, of which ∼496 million were retained after quality filtering and demultiplexing (mean = 2.2 million reads per individual). Initial variant calling generated ∼1 million variant sites, which reduced to 38,615 loci after stringent filtering. Data from all 222 individuals were retained as coverage was relatively high (mean = 9.5×; ∼1 locus per 13kb based on genome size of ∼528 Mb [B. Richardson, pers. comm.]), and individuals had relatively low percentages of missing data (mean = 5.9%).

While multilocus *F*_ST_ estimates were in the low range (0.076; 95% bootstrapped CI: 0.067-0.085; see Fig. S3 for population pairwise estimates), genetic structuring among populations was clearly indicated by tight, often non-overlapping population clusters in principal component space, and a strong concordance between geographic and genetic distances (Procrustes correlation coefficient *r* = 0.738; Fig. 1B). Mantel tests of IBD and IBE indicated that both geographic distance and environmental distance predicted genetic distance among populations (IBD: Mantel *r* = 0.246, *P* = 0.025; IBE: Mantel *r* = 0.286, *P* = 0.008). There was less evidence for an association between environmental and geographic distance (Geo-Env: Mantel *r* = 0.202, *P* = 0.084; Fig. 2). Similar results of IBD and IBE were found using the piecewiseSEM approach (IBD: direct β = 0.293, indirect β = 0.086; IBE: β = 0.300; Geo-Env: β = 0.286; Fig. 2D), but relative contributions of each metric could be assessed using this holistic approach. As our piecewiseSEM incorporates multiple response variables and random effects, we could further assess the model through the variance explained (*r*^2^) for each response variable with fixed effects only (marginal estimation) and with random effects (population) included (conditional estimation). Thus, variance explained was relatively low for the marginal estimation with only fixed effects (environmental distance: *r*^2^ = 0.075, genetic distance: *r*^2^ = 0.186), but was much greater in the conditional estimation on the full model including random effects (environmental distance: *r*^2^ = 0.744, genetic distance: *r*^2^ = 0.403). This indicated that including a random effect of population identification to eliminate the inherent autocorrelated structure of pairwise comparisons was most appropriate. Additionally, populations displayed moderate levels of genetic diversity (θ*_π_* = 0.019–0.021; Table 1) and there was no evidence of selfing/inbreeding as individual inbreeding estimates, *F*, were effectively 0 (mean = 0.0033; only one individual with *F* > 0.035).

**Figure 2.**
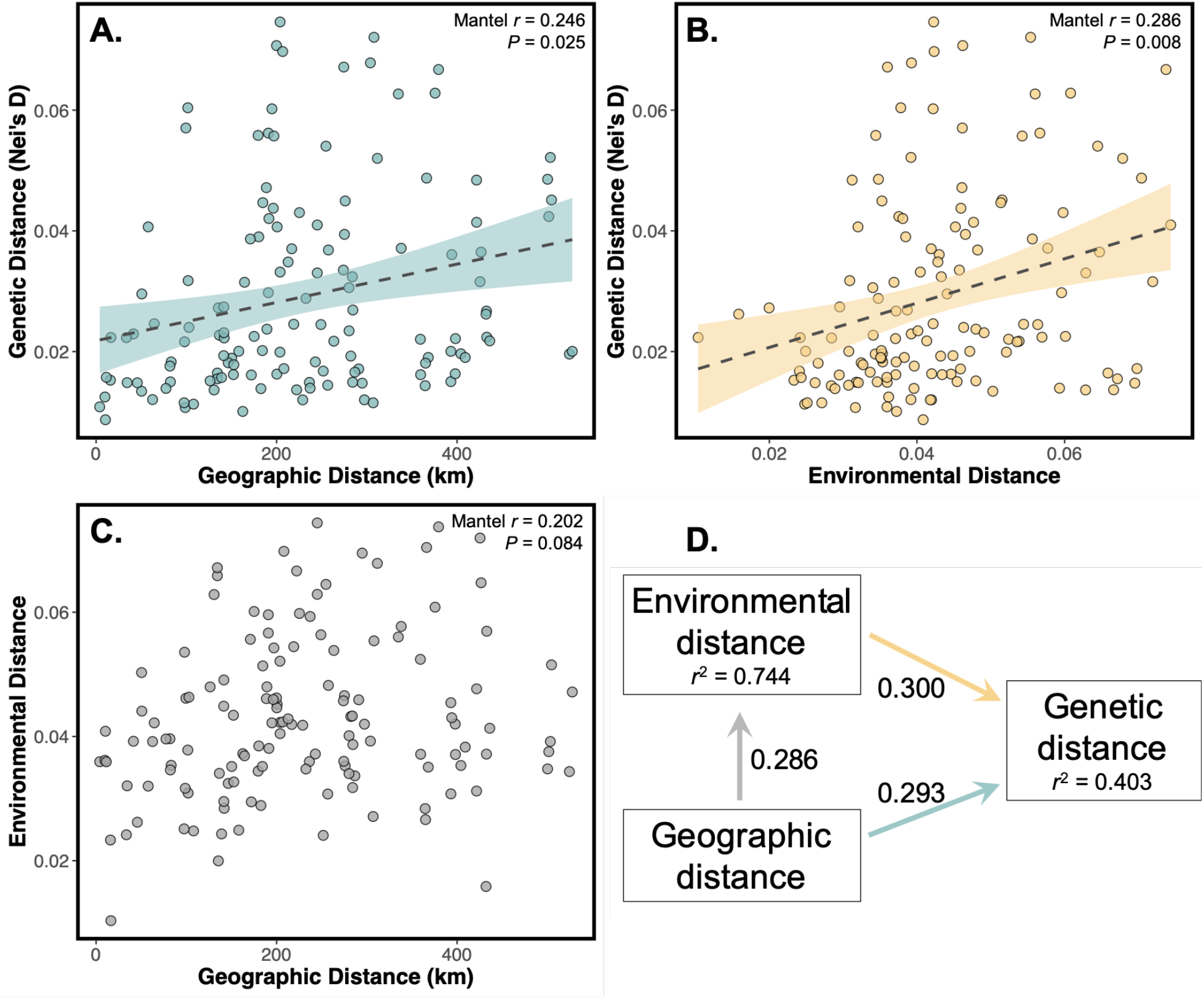
Neutral and adaptive processes shape landscape genomic structure. Positive relationships between genetic distance and both (**A**) geographic and (**B**) environmental distances provide evidence for IBD and IBE. (**C**) Less evidence for an association between geographic and environmental distance is most likely due to the heterogeneous environments of the Great Basin. (**D**) Relative contributions of environmental and geographic distance to genetic distance were estimated using a piecewiseSEM. Geographic distance had a slightly larger relative impact on genetic distance (β = 0.379; or 0.293 + [0.286 × 0.300]) than on environmental distance when accounting for indirect effects. Conditional estimates (includes random effects) of variance explained are represented by the *r*^2^ under each response variable.

**Table 1:** Relevant environmental information for the sampled populations, and summary statistics for sampling effort and genetic diversity (θ_π_). Inbreeding coefficients (*F*) are not reported, as most are effectively zero (mean = 0.0033). Grey rows indicate populations included in genetic analyses but not in the common garden.

| Site Name, State | Site abbr. | Precip. seasonality | Annual mean temp. (°C) | Aspect | Elev. (m) | N | $\theta_\pi$ |
| --- | --- | --- | --- | --- | --- | --- | --- |
| Austin Summit, NV | AS | 0.322 | 7.72 | 244.80 | 2408 | 15 | 0.0206 |
| Bald Mountain, NV | BM | 0.191 | 6.85 | 158.20 | 2245 | 14 | 0.0211 |
| Buena Vista, OR | BV | 0.286 | 8.02 | 127.88 | 1277 | 15 | 0.0209 |
| Dayton Hill, NV | DH | 0.600 | 10.54 | 342.90 | 1400 | 15 | 0.0210 |
| Diamond Crater, OR | DC | 0.267 | 8.01 | 138.37 | 1295 | 14 | 0.0211 |
| East Walker, CA | EW | 0.531 | 8.15 | 82.69 | 2043 | 10 | 0.0195 |
| Finger Rock, NV | FR | 0.185 | 8.45 | 9.46 | 2129 | 15 | 0.0205 |
| Hwy 140, NV | HO | 0.322 | 9.01 | 135.00 | 1423 | 13 | 0.0212 |
| Jones Canyon, NV | JC | 0.406 | 9.25 | 205.02 | 1440 | 15 | 0.0209 |
| Long Valley, CA | LV | 0.579 | 9.02 | 37.41 | 1660 | 12 | 0.0194 |
| Modoc, CA | MD | 0.403 | 8.51 | 315.00 | 1333 | 15 | 0.0202 |
| Patagonia, NV | PT | 0.469 | 10.72 | 24.78 | 1421 | 7 | 0.0204 |
| Peavine Low, NV | PL | 0.643 | 9.09 | 38.66 | 1724 | 14 | 0.0205 |
| Petrified Wash, NV | PW | 0.186 | 9.03 | 266.99 | 1871 | 10 | 0.0213 |
| Smith Creek, NV | SC | 0.175 | 7.73 | 170.75 | 2050 | 14 | 0.0202 |
| Spanish Springs, CA | SS | 0.307 | 7.54 | 356.19 | 1645 | 9 | 0.0195 |
| Virginia Mountains, NV | VM | 0.561 | 9.64 | 145.01 | 1503 | 15 | 0.0209 |

### Genetic-environment association analyses

Genetic-environment association analyses provided evidence of local adaptation to specific environmental variables and an overlapping set of loci potentially under selection was detected by the two methods employed. The signature of local adaptation could be attributed to a set of environmental predictors that were largely consistent between the two methods employed (Table 2). The first RDA axis identified 176 outlier loci potentially associated with adaptation (*PVE* = 21.65%; Fig. 3B). The environmental variables with the greatest explained variance in the RDA included precipitation seasonality, slope, elevation, and annual precipitation (Table 2; Fig. 3). Gradient Forest results overlapped considerably with those from the RDA, with loadings from the first RDA axis and *R*^2^ from GF being strongly correlated for both the environmental predictors and the identified outlier loci (environment: Pearson’s *r* = 0.796, loci: Pearson’s *r* = 0.656; Fig. 4). Precipitation seasonality was overwhelmingly the strongest environmental predictor in both the RDA and GF analyses, with both the largest explained variance and number of associated outlier loci (56.5% of the total outliers). Consistent sets of loci were identified as outliers in both analyses: 54.5% (96/176 loci) had allele frequency shifts associated with environmental variation in the RDA and as estimated by split importance (or *R*^2^ *>* 0) in GF. A permutation test (*n* = 1000) using 176 randomly sampled loci across all 38,615 loci indicated the RDA outlier loci had more overlapping loci with GF outliers than would be expected by chance (permuted outlier range: 3.4–17.0% [6–30 loci]; *P <* 0.001). Furthermore, 65 out of the 96 RDA and GF outlier loci were most strongly associated with the same environmental predictor. An additional 27 loci (92 or 95.8% of RDA and GF outlier loci) had the same top two environmental associations, with minimal difference (average *<* 0.095 SD) between the first and second association strength to that locus (*r* for RDA or *R*^2^ for GF). In other words, RDA and GF approaches typically agreed on the top two environmental predictors driving the outlier loci, only flipping between first and second when the difference in association strength was negligible.

**Table 2:** Environmental variable abbreviations with descriptions and units used in the text and figures, and the range of values for each variable. Data for these variables are represented more fully in Table S3. Both genetic and phenotypic associations with each environmental variable are given for RDA (loading on the first axis), GF (split importance *R*^2^), and RF (importance *R*^2^). Rank represents the rank of that variable/analysis combination (i.e., 1 is most important, 10 is least). Number of loci is the number most highly correlated with each environmental variable with percent (%) out of the total outlier loci overlapping between RDA and GF (92 loci). Variables are ordered by their loading on the first RDA axis.

| Env. variable | Description (units) | Variable range | Genetic-environment association |  |  | Common garden |
| --- | --- | --- | --- | --- | --- | --- |
| | | | RDA1 (rank) | GF $R^2$ (rank) | # loci (%) | RF $R^2$ (rank) |
| PrcpSeas | Precipitation seasonality<sup>1</sup> | 0.18-0.64 | 0.666 (1) | 0.026 (1) | 52 (56.5%) | 20.82 (1) |
| Slope | Slope (degrees) | 0.68-17.39 | 0.538 (2) | 0.012 (5) | 6 (6.5%) | 1.15 (8) |
| Elevation | Elevation (m) | 1277-2408 | 0.526 (3) | 0.014 (2) | 13 (14.1%) | 4.91 (7) |
| MinVPD | min. vapor pressure deficit (hPa) | 1.28-3.30 | 0.497 (4) | 0.009 (6) | 3 (3.3%) | 6.59 (6) |
| PrcpAnn | Annual precip. (mm) | 215.19-388.44 | 0.431 (5) | 0.014 (3) | 1 (1.1%) | 6.69 (5) |
| Aspect | Aspect (degrees from North) | 9.46-356.19 | 0.409 (6) | 0.012 (4) | 11 (12%) | 12.31 (2) |
| SoilMax | max. soil water capacity (cm) | 89.77-163.34 | 0.362 (7) | 0.007 (10) | 1 (1.1%) | 0.00 (9; tie) |
| CumlAET | Cuml. actual evapotranspiration (mm per year) | 185.29-275.46 | 0.299 (8) | 0.008 (9) | 4 (4.3%) | 0.00 (9; tie) |
| TempAnn | Annual temp. (°C) | 7.1-10.7 | 0.267 (9) | 0.008 (8) | 1 (1.1%) | 10.75 (3) |
| HL | Heat load index<sup>2</sup> | 0.81-1.03 | 0.24 (10) | 0.008 (7) | NA | 9.87 (4) |

1. PrcpSeas classes designated by Walsh & Lawler, 1981
2. HL calculated from McCune & Keon, 2002

**Figure 3:**
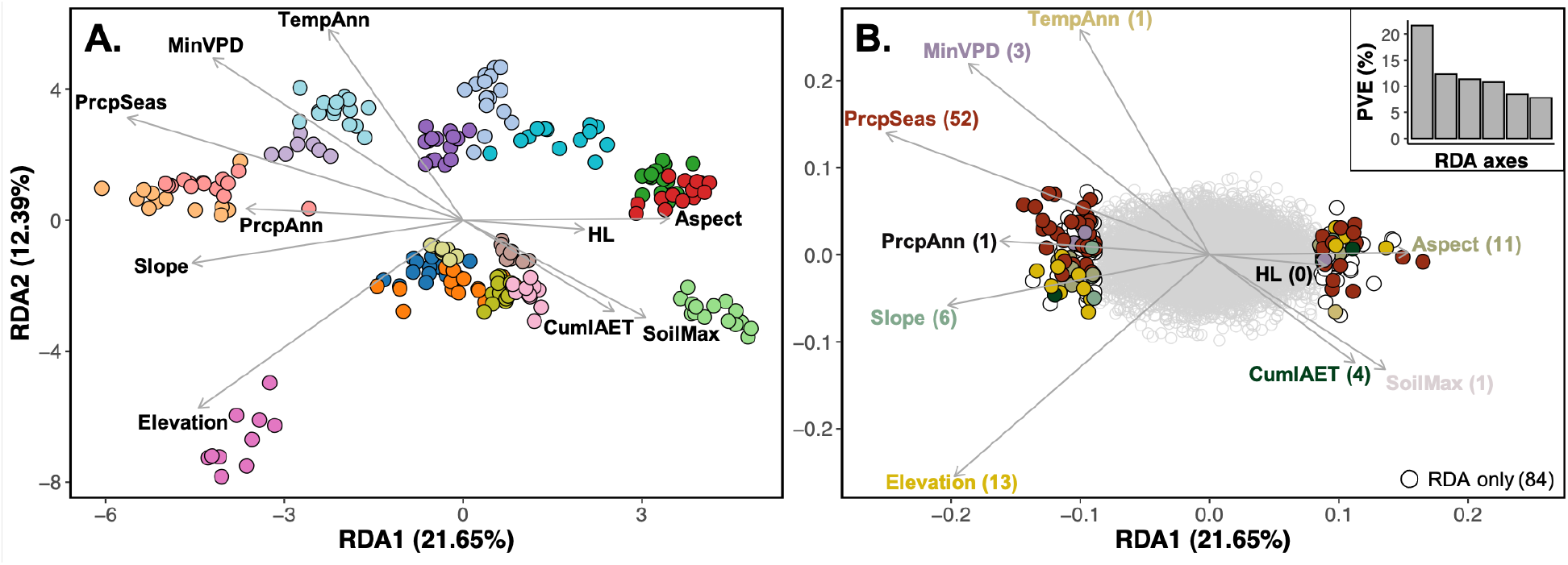
Environmental variation predicted population structure and was associated with pronounced allele frequency shifts at loci likely to be under selection. (**A**) Redundancy analysis (RDA) of the genotype probabilities for each individual associated with environmental predictor variables (Table 2). Direction and length of arrows correspond to the loadings of each variable on the same two RDA axes. Point colors correspond to population colors in Fig. 1. (**B**) The loadings of each locus onto the same two RDA axes and environmental predictor variables (vectors scaled 8.45× and 0.376× for graphical representation in A and B, respectively). The colored points indicate loci identified as outliers (*±*3.5 SD; *P <* 0.0005) on the first RDA axis. The top-right inset illustrates the proportion of variance explained (*PVE*) for each of the first six RDA axes, with the first RDA axis overwhelmingly explaining the most variance. Points are colored to match the environmental predictor of greatest association in both RDA and GF analyses with white representing outliers unique to RDA. The number of outlier loci associated with each variable is represented parenthetically. Environmental correlations can be positive or negative, thus the same colors can exist for outliers on the left and right.

**Figure 4:**
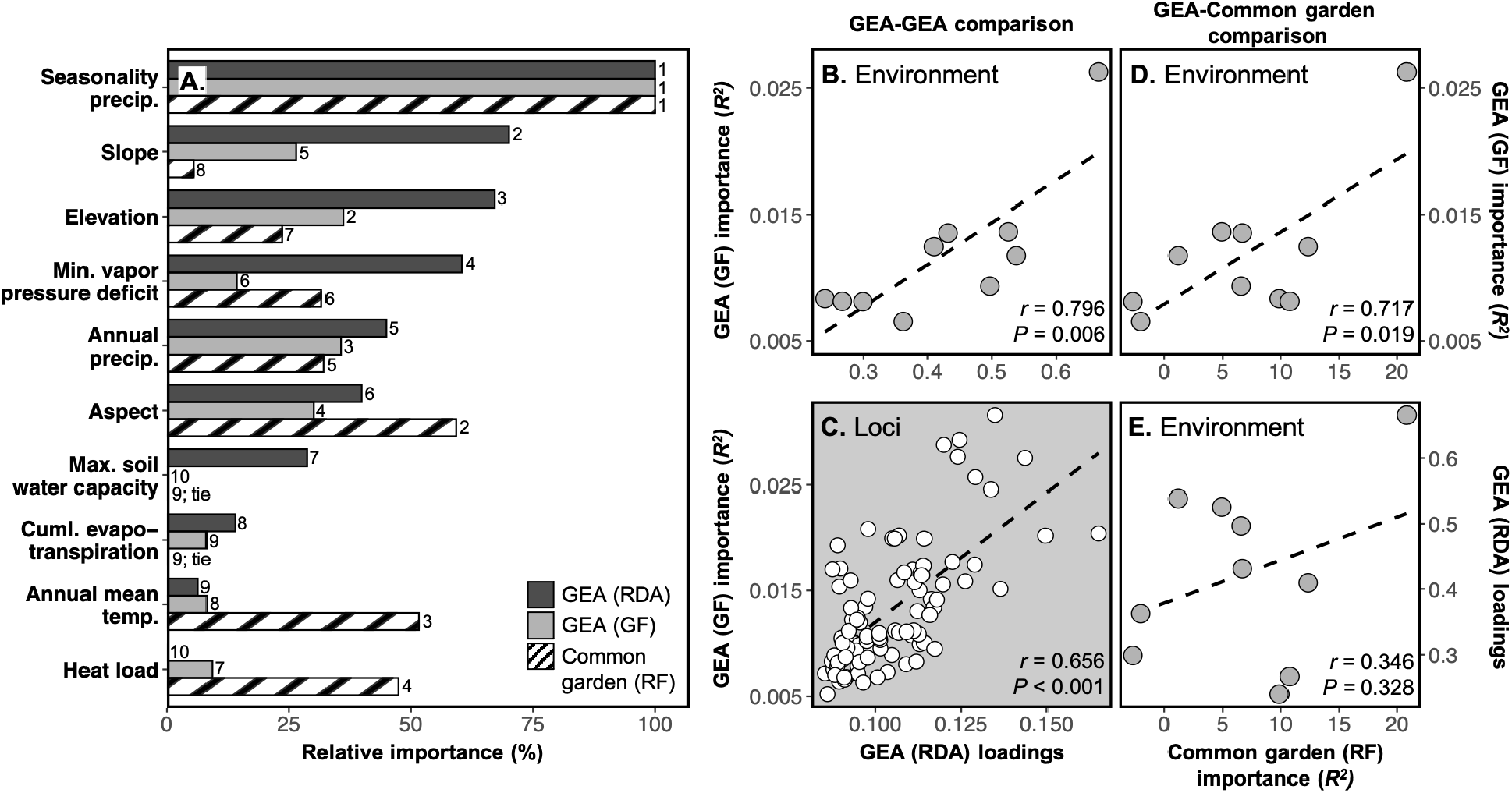
Local adaptation to specific environmental predictors illustrated across both GEA approaches and analyses of phenotypic data from a common garden. (**A**) Relative importance of each environmental variable (scaled from 0 to 100%, with 100% having the largest contribution) for both the phenotype-environment and genetic-environment associations. GEA analyses are represented in solid colors with loadings of the first RDA axis (dark) and split importance (*R*^2^) from GF (light). Relative importance of each environmental variable in the phenotypic analysis is represented by the importance *R*^2^ from RF and represented by the striped bar. The composite variable of seedling emergence, seed weight, and shoot biomass was used in the RF analysis to represent phenotypic variance within the common garden. The rank of each environmental variable within its analysis is displayed to the right of each bar. Concordance among the two GEA approaches is illustrated by strong correlation between the absolute value of the first RDA axis loadings and GF *R*^2^ for each environmental variable (**B**) as well as across the individual outlier loci (**C**). Concordance of environment variable association in the common garden with both GEA analyses is displayed in panels **D** and **E**.

### Phenotypic variation

Populations grown from wild collected seed in the common garden differed in all measured phenotypes (seedling emergence: *P <* 0.001; seed weight: *P <* 0.001; days to emergence: *P <* 0.001; shoot biomass: *P* < 0.001; average root diameter: *P* = 0.014; Figs. 5 and S4). Variation among populations was high; for example, mean seed weight and shoot biomass were 3.3× and 4.3× greater, respectively, in the largest populations over the smallest (Fig. 5). Correlations among environmental and phenotypic variables were consistent with local adaptation, as phenotypic variation was associated with environmental variation across populations (Figs. S1 and S2). Only seedling emergence, seed weight, and shoot biomass had positive variance explained in the RF analyses with precipitation seasonality as overwhelmingly the largest predictor of phenotypic variation among populations (seedling emergence: *r* = 0.790, seed weight: *r* = 0.710, shoot biomass: *r* = 0.635; Figs. 4 and 5). Days to emergence and average root diameter, while each having positive correlations to specific environment variables (Figs. S1 and S2), had negative variance explained in RF indicating that broadly, when all ten environmental variables are included, environmental variables are no better at predicting variation in those two phenotypes than random chance.

**Figure 5:**
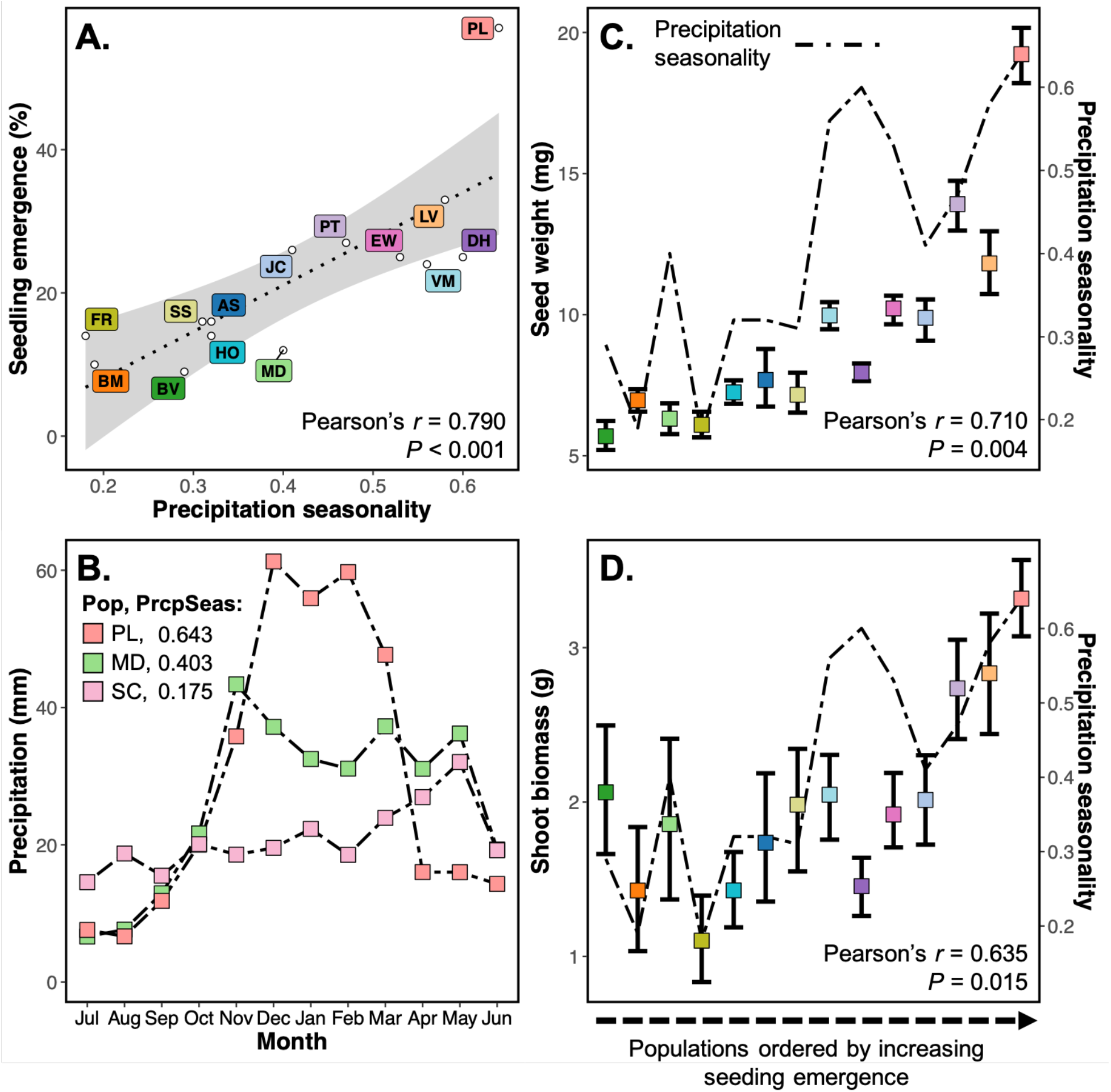
Phenotypic data from a common garden suggest local adaptation to precipitation seasonality. (**A**) Strong associations between seedling emergence and precipitation seasonality suggests local adaptation. Seedling emergence is calculated as the percentage of plants emerged and survived 40 days in each population. (**B**) Monthly precipitation averages from 30-year PRISM normals for three populations with the highest (PL), lowest (SC), and median (MD) precipitation seasonality. (**C**) and (**D**) Seed weight and shoot biomass vary by orders of magnitude among the populations. Additionally, their associations with precipitation seasonality (described by Pearson’s *r* and *P*-values) are consistent with local adaptation. The line corresponds to the precipitation seasonality estimate of that population. The mean phenotype and 95% bootstrapped confidence intervals are represented by the squares and error bars. The populations are ordered by seedling emergence, illustrating correlation among the phenotypes. Colors are the same as those in the Fig. 1 map.

*P*_ST_ - *F*_ST_ comparisons indicated seedling emergence, seed weight, and shoot biomass were more differentiated than neutral expectation (i.e., directional selection: *P*_ST_ > *F*_ST_) while days to emergence and average diameter were not (Table S4, Fig. S5). Furthermore, the critical values of the *c/h*^2^ ratio for seedling emergence, seed weight, and shoot biomass were low (0.042 and 0.578, respectively) indicating the robustness of our comparisons and conclusions. Because days to emergence and average diameter were not predicted by environmental variation and did not deviate from neutral expectations, the composite variable of seedling emergence, seed weight, and shoot biomass was used as representative of phenotypic variation when assessing environmental associations and comparing to the GEA analyses (Table 2). There was considerable overlap in the environmental variables inferred to underlie local adaptation in both GEA (genetic) and common garden (phenotypic) analyses. Exceptions include annual mean temperature and heat load, which were the third and fourth-ranked predictors of phenotypic variation but were relatively insignificant in the GEA analyses (Fig. 4A). Even still, variable importance values for environmental variables were strongly correlated between the GEA and common garden (RF ∼ GF: *r* = 0.717, RF ∼ RDA: *r* = 0.346), mostly driven by associations with precipitation seasonality in both analyses.

Further evidence for the importance of environmental and genetic variation on the phenotypic variation was provided by the variance partitioning approach combined with RDA and pRDAs. Taken together, genetics, environment, and geography explained a large proportion of phenotypic variance (*PVE* = 69.84%; Table 3, Fig. S6, but when each variable was assessed individually, environment (*PVE* = 54.50%) and genetics (*PVE* = 52.66%) explained substantially more variance than geography (*PVE* = 21.26%). When looking at the partial contributions of each predictor independent of correlative effects of other predictors, environment contributed the most (adj. *r*^2^ = 16.12%), followed by genetics (adj. *r*^2^ = 12.94%), and geography contributing very little (adj. *r*^2^ = 2.57%).

**Table 3:** Variance partitioning of the individual and shared contributions of genetic, environmental, and geographic variation on phenotypic variation from the common garden experiment. All three predictors captured 69.84% of the total phenotypic variation. Adjusted *r*^2^ represents the individual contribution of the predictor with all others partialled out and the proportion of variance explained (*PVE*) represents the overall contribution without controlling for interactive effects among the predictors. Models are ordered by the greatest *PVE*.

| Model | Adj. $r^2$ (%) | $PVE$ (%) |
| --- | --- | --- |
| Pheno ~ Gen + Env + Geo | 20.36<sup>–</sup> | 69.84<sup>*</sup> |
| Pheno ~ Gen + Env | 19.52<sup>–</sup> | 67.28<sup>*</sup> |
| Pheno ~ Env + Geo | 0.00<sup>–</sup> | 56.90<sup>*</sup> |
| Pheno ~ Env | 16.12<sup>*</sup> | 54.50<sup>*</sup> |
| Pheno ~ Gen + Geo | 0.00<sup>–</sup> | 53.72<sup>*</sup> |
| Pheno ~ Gen | 12.94<sup>*</sup> | 52.66<sup>*</sup> |
| Pheno ~ Geo | 2.57<sup>*</sup> | 21.26<sup>*</sup> |
| Residuals | 30.16<sup>–</sup> | – |

\* model significance ( $P \leq 0.05$ ) using RDA/pRDA – model significance cannot be tested

## Discussion

An understanding of the distribution and function of landscape genomic variation in widespread, foundational species can provide insight into the evolution and maintenance of diversity and can be critical for managers seeking to preserve or restore communities facing climate change or other disturbances. Although phenotype-environment and genetic-environment association analyses have each been widely employed, relatively few studies have applied both approaches, although their combined application could provide more comprehensive inference of local adaptation (Breed et al., 2019; Rellstab et al., 2015; Sork et al., 2013). Studies that have applied both approaches often report congruent evidence for the environmental contributions to local adaptation (e.g., Carvalho et al., 2020; De Kort et al., 2014; de Villemereuil et al., 2018; Mahony et al., 2020; Yoder et al., 2014). Consistent with such studies, our phenotype-environment and genetic-environment association analyses generally implicated a consistent set of environmental variables and provided complementary evidence for local adaptation across the western Great Basin. While it can be challenging to separate the effects of maternal environment and genetics on trait variation in wild plants, the congruent environmental results from GEA analyses support the inference that phenotypic differentiation in the common garden was at least partially a genetically-based result of local adaptation.

Genetic-environment association analyses have increasingly been applied to studies of local adaptation because they can assess large numbers of individuals and populations across wide environmental gradients and can be readily applied in taxa where common garden designs are not feasible (e.g., endangered species, species that are challenging to grow or have high seed dormancy, are long-lived, or highly vagile). However, GEA analyses can be limited by the spatial resolution of environmental data (Daly, 2006), confounding signals from drift or other aspects of population history (Hoban et al., 2016; Lotterhos & Whitlock, 2014), and potentially weak and diffuse signal underlying polygenic adaptation (Barghi et al., 2020; Rockman, 2012). Common garden approaches have drawbacks as well, including the challenges of reducing environmentally mediated maternal effects, limited phenotypic variation based on sampling or incomplete seed germination, laborious execution, unintended selection imposed by the garden itself, and the exclusion of important ecological processes (e.g., herbivory or competition; Antonovics & Primack, 1982; Gibson et al., 2016; Sork et al., 2013). Additionally, common gardens tend to focus on phenotypes that are suspected to be components of fitness, but that might vary in the direction and magnitude of their effect on fitness in particular environments (Sork et al., 2013). While GEA analyses have been independently applied in restoration contexts without consideration of phenotypic data (e.g., Massatti et al., 2018, 2020; Shryock et al., 2017, 2021), studies utilizing both approaches should provide a more detailed understanding of how both genetic and phenotypic are variation shaped by adaptation to environments. Below we discuss genomic and phenotypic variation of *E. nauseosa* in the context of environmental heterogeneity across the western Great Basin, the distribution and causes of local adaptation, and the potential for such analyses to inform restoration.

### Spatial genomic structure

The degree of fine-scale spatial genomic structure we detected was surprising given *E. nauseosa* is wind-dispersed and fairly continuously distributed across its range (Anderson, 1984). Although levels of genetic differentiation between populations were by no means high (multilocus *F*_ST_ = 0.076; see Fig. S3 for pairwise population estimates), genetic differentiation across fine geographic scales was illustrated by tight and often non-overlapping clusters in PCA space. For example, several pairs of sampling localities separated by < 10 kilometers (e.g., FR to PW; LV to PL; BV to DC) formed wholly distinct clusters (Fig. 1B). The number of loci generated with our sequencing design likely improved our ability to recover such structure, which also appears to be influenced by the environmental and topographic complexity of the western Great Basin. Out-crossing appears to dominate in these populations, as individual inbreeding coefficient (*F*) estimates were close to 0 (mean *F* = 0.003). Although past studies of *Ericameria* have suggested high rates of self-fertilization (Anderson, 1966; McArthur et al., 1978), low levels of inbreeding indicated by our analyses could result from our sampling of adult plants if self-fertilization or inbreeding were associated with reduced seed set or seedling fitness. Further, estimates of nucleotide diversity (θ*_π_* = 0.019–0.021; Table 1) revealed moderately high and consistent levels of standing variation within the populations spanning our study region.

Genetic differentiation among populations was predicted by both geographic and environmental distances (Figs. 1B and 2), indicating that population genetic structure is driven by stochastic processes, such as genetic drift, as well as those stemming from environmental variation. A positive relationship between environmental and genetic distances among populations can arise from IBE, where gene flow is reduced across environmental gradients such as when divergent selection reduces effective migration among populations adapted to different local conditions (Shafer & Wolf, 2013; Wang & Bradburd, 2014). Across our sampled populations, environmental and geographic distances were weakly related (Fig. 2C), eliminating some of the confounding demographic processes shaping spatial genomic variation that can potentially interfere with the detection of local adaptation. Similar studies of heterogeneous regions of western North America have gone as far as estimating the relationship between geography and environment prior to sampling to ensure diverse environmental ranges and reduce confounding effects for inferring genetic-environment associations (e.g., Massatti et al., 2018; Massatti & Knowles, 2020). Here, we used a holistic approach (piecewiseSEM) to estimate the relative effects of geography and environment on genetic structure while accounting for spatial autocorrelation (Fig. 2D). Environment had a slightly larger direct influence than geography on genetic distance among populations. Results from additional analyses, including constrained ordination (RDA) and machine-learning approaches (GF), are also consistent with an influence of environment on spatial genomic structure. Overall, our results suggest that local adaptation and reduced gene flow across environmental gradients has contributed to spatial genetic structure, and that geographic proximity alone is not a great predictor of the environmental conditions underlying local adaptation in this region (Hereford, 2009; Raabová et al., 2007). Thus, our results support the development of spatially-complex seed transfer zones (areas where seeds can be moved without loss of fitness) that are based on environmental, rather than geographic, distances (e.g., Bradley St. Clair et al., 2013; Massatti et al., 2020).

### Genetic-environment associations

While reduced representation sequencing data have limits for characterizing the genetic basis of adaptation (Lowry et al., 2017; but see Catchen et al., 2017; McKinney et al., 2017), our sequencing approach generated moderate marker density data (38,615 loci across the estimated ∼528 Mb genome size or ∼1 locus per 13 Kb), providing a sampling of genomic variation likely shaped by a continuum of evolutionary processes. Genetic-environment analyses have advantages compared to more widely employed divergence outlier scans (i.e., *F*_ST_ outlier analysis) for inferring adaptation as they interrogate environmental variation associated with allele frequency shifts while detecting locus-specific signals of selection (Capblancq et al., 2018; Forester et al., 2016, 2018; Lotterhos & Whitlock, 2015; Rellstab et al., 2015). The two GEA methods we employed (RDA and GF) detected an overlapping set of loci tagging genomic regions likely responding to local environmental variation (Figs. 3 and 4C). The RDA detected 176 outlier loci (0.4% of total loci) associated with environmental variation, 96 of which were dually implicated by GF and largely attributed to the same environmental influence (95.8%; 92 out of 96 loci). While GF has largely been used for genomic prediction of maladaptation under future climate change scenarios (e.g., Bay et al., 2018; Capblancq et al., 2020; Keller et al., 2018), Fitzpatrick et al., (2021) recently showed that it also performs well for outlier loci detection. Combined and congruent results from RDA and GF provide genomic evidence for local adaptation of *E. nauseosa* populations across environmental space in the western Great Basin, even in the absence of phenotypic information.

The two GEA approaches also suggested a congruent set of environmental variables have influenced local adaptation, with the strongest contributions from precipitation seasonality, slope, aspect, elevation, and annual precipitation (Table 2, Fig. 4A,B). Although the relative contribution and order of importance varied for genetic-environment association across methods, the influence of environmental variables and their association with outlier loci were strongly correlated (Fig. 4B,C). Many of the implicated environmental variables are known to predict spatial variation in plant performance (Barga et al., 2018; Dilts et al., 2015; Stephenson, 1998) and are often thought to influence natural selection across plant species ranges (Dawson et al., 2000; Loveless & Hamrick, 1984; Reich et al., 2003). For example, in a meta-analysis of 161 common garden-based studies in the Great Basin, 75% identified locally adapted traits associated with environmental variables similar to those implicated here (e.g., seasonality, temperature, precipitation, soil, evapotranspiration, etc.; Baughman et al., 2019). Precipitation seasonality, a metric also strongly correlated with fall actual evapotranspiration (*r* = 0.824), was overwhelmingly the strongest predictor in our analyses (Figs. 3 and 4). This variable has similarly been inferred to influence local adaptation in other recent GEA analyses of arid land plants (Shryock et al., 2017; Temunović et al., 2020). Precipitation seasonality values for our sampled localities ranged from 0.175 – 0.643 (Tables 1 and 2), with locations having higher seasonality experiencing relatively high precipitation in the winter months (Fig. 5B; Walsh & Lawler, 1981). *Ericameria nauseosa* is a phreatophyte that develops a deep root system to access soil moisture and avoid low water potentials in the summer, when precipitation is scarce (Donovan & Ehleringer, 1994). The importance of precipitation seasonality for predicting phenotypic variation, including seedling emergence and seed size, suggest that plants from more seasonal environments could be avoiding summer drought by relying on deep stores of water from winter precipitation (Branson et al., 1976). Additional variables implicated by GEA analyses include slope, aspect, elevation, and annual precipitation, all factors that influence soil depth, water table level, and available water during summer drought (Måren et al., 2015; Van de Water et al., 2002).

### Phenotypic variation in a common garden

Common garden experiments have long been used to quantify genetically-based phenotypic differentiation among populations and to infer local adaptation to environment (e.g., Baughman et al., 2019; Kawecki & Ebert, 2004; Lind et al., 2018). In our study, seed and seedling traits were substantially differentiated among populations (Figs. 5 and S4), suggesting they are controlled to some degree by genetic variation among populations. Further, phenotypic and environmental variation were strongly associated across sites, suggesting local adaptation driven by specific environmental variables (Figs. 4, 5, S1, and S2). Phenotypic variation among populations for traits exhibiting environmental associations was also greater than expected under drift after accounting for population structure (i.e., *P*_ST_ > *F*_ST_; Table S4, Fig. S5). However, because we grew and measured seedlings from wild collected seeds, our design does not account for maternal effects (Gienapp et al., 2008; Kawecki & Ebert, 2004). Environmental variation could shape maternal effects through, among other mechanisms, proximate influences on seed size (Donohue, 2009; Lampei et al., 2017; Roach & Wulff, 1987; Sultan et al., 2009), and *E. nauseosa* seed size was correlated with environmental variation (e.g., precipitation seasonality and annual precipitation; Fig. S1, S2) as well as other measured phenotypes (e.g., seedling emergence and shoot biomass). Thus, phenotypic differentiation could have arisen from genetic variation, maternal effects, or a combination the two. Accounting for maternal effects can be challenging, requiring multiple generations of seed production in common gardens, and may not be feasible for many long-lived arid land plants or over management relevant timeframes. Indeed, a recent review of phenotype-environment association analyses of Great Basin plants found that only 19 (5.8%) of 327 experiments accounted for maternal effects (Baughman et al., 2019). While we cannot easily distinguish among proximate and ultimate causes of seed weight variation across environments, our results are nonetheless consistent with past studies demonstrating that germination, emergence phenology, and seed and seedling traits are often heritable and evolve in response to environmental variation (e.g., Baughman et al., 2019; Caignard et al., 2019; Clauss & Venable, 2000; Gremer & Venable, 2014; Hernández et al., 2019).

Congruence in the environmental variables implicated in phenotype-environment and genetic-environment association analyses also support the inference that phenotypic differences measured in the common garden are influenced by local adaptation. Consistent with GEA analyses, precipitation seasonality had the strongest influence on seed and seedling phenotypic variation. Increases in this variable were associated with increased seedling emergence, seed weight, and shoot biomass (Figs. 5 and S1), possibly because the concentration of precipitation in the winter favors larger seeds that are more likely to germinate and produce larger seedlings (Brown et al., 1997). While aspect, annual mean temperature, and heat load also predicted phenotypic variation (Figs. 4 and S1), the specific timing and magnitude of precipitation was more predictive. Increased annual mean temperature and precipitation were also associated with greater seedling emergence, heavier seeds, and bigger shoots (Figs. S1 and S2). Temperature is known to influence germination and subsequent emergence in *E. nauseosa*: seeds from warmer habitats germinate more rapidly, while those from colder montane environments have greater dormancy and require extended periods of cold-wet stratification (Meyer et al., 1989). Finally, increasing soil water capacity was associated with thicker and shorter roots (Figs. S1, S2, and S4). While these soils have a greater capacity to store water, their higher clay content and finer texture lead to lower water potentials and greater water limitation during summer. Thus, thicker roots may increase fitness by reducing root cavitation in these soils (Sperry & Hacke, 2002).

While inferences were broadly congruent between the phenotype-environment and genetic-environment association analyses, there were some inconsistencies as well. For example, annual mean temperature was strongly associated with phenotypic variation but was a minor contributor to GEA. Similarly, slope and elevation ranked highly in the GEA analyses but low in the phenotype-environment association analyses (Fig. 4). Several factors, both within our experiment specifically and more generally, could have contributed to this discordance. First, our common garden focused on emergence phenology and seedling phenotypes suspected to have fitness consequences early in life, while the GEA analyses were based on variation in adult plants that survived in the wild for multiple years. Second, discordance may have arisen from the selective environment of the common garden itself (Gibson et al., 2016). That is, because not every sown seed germinated and survived, the common garden may have unintentionally represented a non-random set of phenotypic variation, which is almost inevitable when growing wild plants. Finally, strong phenotype-environment associations without strong GEA support could arise from traits more strongly influenced by maternal environment. Thus, the incomplete congruence among inferences of local adaptation from phenotype-environment could represent fruitful areas for further study.

Since we sampled genetic variation from adult plants in nature and not from common garden seedlings, we could not directly quantify genetic variation underlying phenotype in the garden (i.e., using genome-wide association). Nonetheless, a variance partitioning approach allowed us to examine the contribution of genetic, environmental, and geographic variation among populations to phenotypic variation measured in the common garden. Genetic and environmental variation were much stronger predictors of phenotypic variation than geography (Table 3, Fig. S6). Environment explained 54.50% of the variance in phenotype alone, while all three variables together explained 69.84% of the variation. As plants were grown and phenotyped in a common garden, the contribution of environment in these analyses likely represents a combination of genetic variation controlling phenotype (i.e., local adaptation to environmental variation across the sampled populations) and maternal effects, while the contribution of genetic variation represents genome-wide differentiation among populations. As many plants are adapted to local environmental conditions (Leimu & Fischer, 2008), these results illustrate the importance of considering environmental variation in heterogenous landscapes where geographic proximity alone may be a poor predictor of local adaptation.

### Implications for Management

Despite being widespread, disturbance oriented, and wind dispersed, *E. nauseosa* of the western Great Basin exhibited fairly pronounced spatial genomic structure, driven both by stochastic processes and local adaptation to environmental variation. Because *E. nauseosa* has not been widely included in large-scale restoration projects to-date, it is an ideal system for examining processes underlying natural genomic variation (i.e., evolutionary history, local adaptation, gene flow, etc.) in plants of the Great Basin as extant populations likely exhibit an unmanipulated representation of genetic diversity across its range. Inference of local adaptation from our population genomic and common garden approaches implicated a largely consistent set of environmental variables and strongly pointed to precipitation seasonality as a major driver of local adaptation. Given the apparent importance of this variable in this and other recent studies of local adaptation in arid land plants (Baughman et al., 2019), seasonality may be generally important for shrub species of the Great Basin and could be incorporated into the estimation of seed transfer zones for taxa that have not been thoroughly studied, in addition to temperature and aridity, which are commonly used for this purpose (e.g., Bower et al., 2014). While there are clear advantages to pairing GEA and common garden approaches for the study of local adaptation (e.g., de Villemereuil et al., 2016), consistent results from both approaches from this and other recent studies (e.g., De Kort et al., 2014; de Villemereuil et al., 2018; Herrera et al., 2015; Yoder et al., 2014) indicate GEA analyses alone can usefully characterize the genetic signature and environmental drivers of local adaptation in organisms where common gardens are not possible or practical. Additionally, our results illustrate how congruence among approaches may bolster inference of local adaptation for phenotypic data measured in common gardens for species where it is difficult to control for maternal effects. While higher marker density data (e.g., whole genome resequencing) will ultimately provide more thorough perspective on the genetic architecture of local adaptation, reduced representation sequencing provides an efficient means to sample large numbers of individuals and environments and can be applied across most organisms. Although such data lacks critical perspective on phenotypic variation and its relation to fitness, it has the additional benefit of providing information on genetic diversity, differentiation, and gene flow across space and environments that could be critical for predicting the response of populations to climate change (Capblancq et al., 2020; Waldvogel et al., 2020) and improving the resolution and accuracy of provenance collections for seed sourcing strategies (Breed et al., 2019; Rossetto et al., 2019).

## Supporting information

SupportingInformationText

## Acknowledgments and conflict of interest statement

This project was funded by grant 2017-67019-26336 from the USDA National Institute of Food and Agriculture Sustainable Agroecosystems program and Cooperative Agreements L16AC00318 and L19AC00013 from the Bureau of Land Management to EAL and TLP. We would like to thank Katie Uckele, Lanie Galland, Joshua Hallas, Bryce Richardson, Christopher Halsch, Thomas Dilts, Andrew Eckert, A. Grant Schissler, and the UNR Plant Insect Group for intellectual, editorial, and analytical guidance during the completion of this project. Thomas Dilts was especially helpful for generating and selecting the environmental variables used in these analyses. We would also like to thank Shannon Swim, Owen Baughman, Meagan O’Farrell, Kalin Ingstad, Sage Ellis, Marenna Disbro, Kelsey O’Neill, Trevor Carter, Amber Durfee, Katelyn Josifko, Carley Crosby, Casey Iwamoto, Madeline Lowe, and Thomas Cramer all of whom assisted in the organization, implementation, and data collection for the common garden experiment conducted at the University of Nevada - Reno greenhouse.

## Data Archiving Statement

The data that supports the finding of this study are available from the Dryad Digital Repository: doi:10.5061/dryad.j0zpc86f4. This includes population-level summaries for phenotypic and environmental data, individual fastq files, filtered VCF file, and a matrix of genotype probabilities. Accompanying scripts for data generation and analyses will be made available after acceptance of the manuscript.

