## SupportingInformationText for "Genomic and common garden approaches yield complementary results for quantifying environmental drivers of local adaptation in rubber rabbitbrush, a foundational Great Basin shrub"

1664 N. Virginia Street

Department of Biology, 100

University of Nevada, Reno

Reno, NV 89557, USA

### Supporting Information Methods

#### 1. Reduced representation sequencing and variant calling

Sequences representing potential contaminants (*E. coli* and PhiX) or various Illumina associated oligos were detected and discarded using a pipeline of Perl and bash scripts (http://github.com/ncgr/tapioca). A custom Perl script was used to correct barcode sequencing errors, trim cut site and barcoded oligo associated bases, and demultiplex reads by individual. Because a reference genome was not available for *E. nauseosa* or a close relative, we used a *de novo* assembly of unique reads to create a consensus reference of genomic regions sampled by our sequencing approach. This reference assembly was generated and optimized using shell scripts and documentation provided for dDocent (Puritz et al., 2014) (cutoffs: individual = 6, coverage = 6; clustering similarity: -c 0.8), utilizing CD-HIT-EST (Fu et al., 2012) for assembly. Demultiplexed reads were mapped to the reference assembly using BOWTIE2 (Langmead & Salzberg, 2012), version 2.3.0, with flags --local --very-sensitive-local.

Sequence variants were identified using SAMTOOLS and BCFTOOLS (Li et al., 2009), version 1.8, and subsequent filtering was completed using VCFTOOLS (Danecek et al., 2011), 0.1.15. We retained only biallelic single nucleotide polymorphisms (SNPs) present in at least 70% of the samples and greater than 50 bp apart if present on the same contig. After additional filtering with custom Python scripts, we retained SNPs with minor allele frequency ≥ 5%, summed depth across samples ≥ 100 or < 10000, and alternate allele call quality ≥ 999. We took several additional steps to ameliorate genotyping bias stemming from the potential misassembly of paralogous genomic regions. Removing loci with excessive coverage and retaining only loci with two alleles present, as above, should ameliorate the influence of misassembled paralogous loci in our data (Hapke & Thiele, 2016; McKinney et al., 2018). Lastly, we retained loci with $F_{IS} > -0.5$

or < 0.5, as misassembly of paralogous genomic regions can lead to abnormal heterozygosity (Hohenlohe et al., 2013; McKinney et al., 2017).

### 2. Genetic diversity and individual inbreeding coefficient estimation

Genetic diversity within populations was quantified by calculating nucleotide diversity ($\theta_{\pi}$; Nei & Li, 1979) for each population using methods that incorporate genotype uncertainty implemented in ANGSD (Korneliussen et al., 2013, 2014). Using individual binary sequence alignment (BAM) files, we estimated the folded site allele frequency likelihoods with "realsfs" and calculated genotype likelihoods using the setting "GL 1" estimated from the model implemented in SAMTOOLS model to obtain the likelihood of the folded site frequency spectrum (SFS). We estimated nucleotide diversity ($\theta_{\pi}$) using "doThetas 1" and "thetastat" commands using SFS likelihoods as priors, respectively, for each locus across the reference assembly and averaged the measures for each population. Additionally, individual inbreeding coefficients (*F*) were calculated while incorporating genotype uncertainty using NGSF (Vieira et al., 2013), implemented through ANGSD-WRAPPER (Durvasula et al., 2016). First, genotype likelihoods were estimated using ANGSD and mapped to the reference assembly with a minimum base quality of 20 and mapping quality of 30. To speed up convergence and account for the inherent low coverage of the data, we use the approximate expectation-maximum (EM) algorithm within NGSF and a minimum of $1e^{-9}$.

### 3. $P_{ST}$ - $F_{ST}$ Comparisons

Assessing the relative contributions of adaptive and neutral processes in shaping phenotypic variation within and among populations can be estimated by comparing quantitative trait and neutral genetic variance (i.e., $Q_{ST}$ - $F_{ST}$ comparisons; McKay & Latta, 2002; Merilä & Crnokrak, 2001). While estimation of $Q_{ST}$ requires breeding programs, an analogous proxy ($P_{ST}$) can be estimated within natural populations or common gardens without pedigrees (Brommer, 2011). The neutral expectation is $P_{ST} = F_{ST}$, and deviations can imply directional ($P_{ST} > F_{ST}$) or stabilizing ($P_{ST} < F_{ST}$) selection. The estimation of $P_{ST}$ is dependent on the $c/h^2$ ratio, where $c$ is additive genetic variance and $h^2$ is narrow-sense heritability. The equation is defined as:

$$P_{ST} = \frac{\frac{c}{h^2}\sigma_B^2}{\frac{c}{h^2}\sigma_B^2 + \sigma_W^2},$$

where $\sigma_B^2$ is the phenotypic variance between populations and $\sigma_W^2$ is the phenotypic variation within populations. Brommer (2011) suggests the null assumption that $c = h^2$, or that the proportion of additive genetic variance is equal for within and between population variances. Because it is known for heritability to vary across populations and the environment it is measured in, additional steps should be taken to ensure robustness of the comparison. We estimated $P_{ST}$ for five phenotypes (seedling emergence, seed weight, days to emergence, shoot biomass, and average diameter) assuming $c = h^2$ with MCMCGLMM in R (Hadfield, 2010) and compared it to our multilocus $F_{ST}$ estimates. Independent linear mixed models were constructed for each phenotype with population as a random effect and block as a fixed effect using distributions of Gaussian for seed weight, shoot biomass, and average root diameter, Binomial for seedling emergence, and Poisson for days to emergence. Each model was run for 100,000 iterations with a 2000 iteration burn-in and sampling every 500 steps. To verify these results were robust across all possible ranges of $c$ and $h^2$ (bounded between 0 and 1), we calculated $P_{ST}$ for each unique $c/h^2$ ratio combination of $c$ and $h^2$ ranging from 0.1 to 1 by a step of 0.1. Additionally, we calculated the critical $c/h^2$ value where the lower 95% CI of $P_{ST}$ equals the upper 95% CI of $F_{ST}$, as suggested by Brommer (2011). The lower the critical $c/h^2$ value indicates greater robustness for the $P_{ST}$ - $F_{ST}$ comparison when $P_{ST} > F_{ST}$ as increasing $c/h^2$ values increase $P_{ST}$ (see Table S5B).

**Supporting Information Literature cited**


Brommer, J. E. (2011). Whither Pst? The approximation of Qst by Pst in evolutionary and conservation biology. *Journal of Evolutionary Biology*, *24*(6), 1160–1168. https://doi.org/10.1111/j.1420-9101.2011.02268.x

Danecek, P., Auton, A., Abecasis, G., Albers, C. A., Banks, E., DePristo, M. A., Handsaker, R. E., Lunter, G., Marth, G. T., Sherry, S. T., McVean, G., & Durbin, R. (2011). The variant call format and VCFtools. *Bioinformatics*, *27*(15), 2156–2158. https://doi.org/10.1093/bioinformatics/btr330

Durvasula, A., Hoffman, P. J., Kent, T. V., Liu, C., Kono, T. J. Y., Morrell, P. L., & Ross-Ibarra, J. (2016). Angsd-Wrapper: Utilities for Analysing Next-Generation Sequencing Data. *Molecular Ecology Resources*, *16*(6), 1449–1454. https://doi.org/10.1111/1755-0998.12578

Fu, L., Niu, B., Zhu, Z., Wu, S., & Li, W. (2012). CD-HIT: accelerated for clustering the next-generation sequencing data. *Bioinformatics*, *28*(23), 3150–3152. https://doi.org/10.1093/bioinformatics/bts565

Hadfield, J. D. (2010). MCMCglmm: MCMC Methods for Multi-Response GLMMs in R. *Journal of Statistical Software*, *33*(2), 1–22. http://www.jstatsoft.org/

Hapke, A., & Thiele, D. (2016). GIbPSs: a toolkit for fast and accurate analyses of genotyping-by-sequencing data without a reference genome. *Molecular Ecology Resources*, *16*(4), 979–990. https://doi.org/10.1111/1755-0998.12510

Hohenlohe, P. A., Day, M. D., Amish, S. J., Miller, M. R., Kamps-Hughes, N., Boyer, M. C., Muhlfeld, C. C., Allendorf, F. W., Johnson, E. A., & Luikart, G. (2013). Genomic patterns of introgression in rainbow and westslope cutthroat trout illuminated by overlapping paired-end RAD sequencing. *Molecular Ecology*, *22*(11), 3002–3013. https://doi.org/10.1111/mec.12239

Korneliussen, T. S., Albrechtsen, A., & Nielsen, R. (2014). ANGSD: Analysis of Next Generation Sequencing Data. *BMC Bioinformatics*, *15*(1), 1–13. https://doi.org/10.1186/s12859-014-0356-4

Korneliussen, T. S., Moltke, I., Albrechtsen, A., & Nielsen, R. (2013). Calculation of Tajima's D and other neutrality test statistics from low depth next-generation sequencing data. *BMC Bioinformatics*, *14*(1), 289. https://doi.org/10.1186/1471-2105-14-289

Langmead, B., & Salzberg, S. L. (2012). Fast gapped-read alignment with Bowtie 2. *Nature Methods*, *9*(4), 357–359. https://doi.org/10.1038/nmeth.1923

Li, H., Handsaker, B., Wysoker, A., Fennell, T., Ruan, J., Homer, N., Marth, G., Abecasis, G., & Durbin, R. (2009). The Sequence Alignment/Map format and SAMtools. *Bioinformatics*, *25*(16), 2078–2079. https://doi.org/10.1093/bioinformatics/btp352

McKay, J. K., & Latta, R. G. (2002). Adaptive population divergence: Markers, QTL and traits. *Trends in Ecology and Evolution*, *17*(6), 285–291. https://doi.org/10.1016/S0169-5347(02)02478-3

McKinney, G. J., Waples, R. K., Pascal, C. E., Seeb, L. W., & Seeb, J. E. (2018). Resolving allele dosage in duplicated loci using genotyping-by-sequencing data: A path forward for population genetic analysis. *Molecular Ecology Resources*, *18*(3), 570–579. https://doi.org/10.1111/1755-0998.12763

McKinney, G. J., Waples, R. K., Seeb, L. W., & Seeb, J. E. (2017). Paralogs are revealed by proportion of heterozygotes and deviations in read ratios in genotyping-by-sequencing data from natural populations. *Molecular Ecology Resources*, *17*(4), 656–669. https://doi.org/10.1111/1755-0998.12613

Merilä, J., & Crnokrak, P. (2001). Comparison of genetic differentiation at marker loci and quantitative traits. *Journal of Evolutionary Biology*, *14*(6), 892–903. https://doi.org/10.1046/j.1420-9101.2001.00348.x

Nei, M., & Li, W. H. (1979). Mathematical model for studying genetic variation in terms of restriction endonucleases. *Proceedings of the National Academy of Sciences of the United States of America*, *76*(10), 5269–5273. https://doi.org/10.1073/pnas.76.10.5269

Puritz, J. B., Hollenbeck, C. M., & Gold, J. R. (2014). dDocent : a RADseq, variant-calling pipeline designed for population genomics of non-model organisms. *PeerJ*, *2*, e431. https://doi.org/10.7717/peerj.431

Vieira, F. G., Fumagalli, M., Albrechtsen, A., & Nielsen, R. (2013). Estimating inbreeding coefficients from NGS data: Impact on genotype calling and allele frequency estimation. *Genome Research*, *23*(11), 1852–1861. https://doi.org/10.1101/gr.157388.113

### Supporting Information Figures

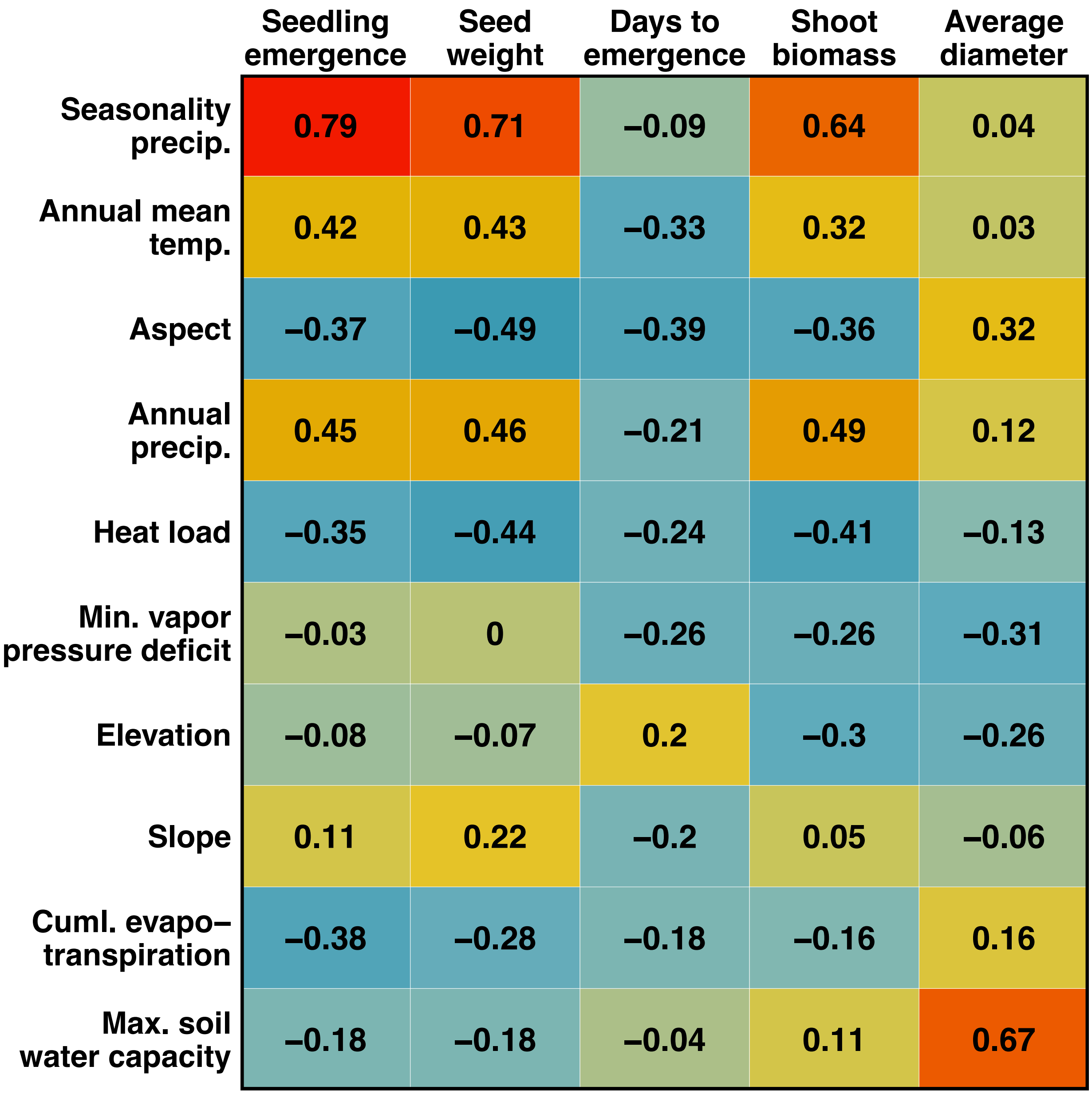


**Figure S1: Heatmap summarizing Pearson correlation coefficients for pairs of environmental and phenotypic variables across populations.** Environmental variables are listed in descending order based on their loading on the first RDA axis. Colors correspond to the magnitude of estimates, with warmer colors corresponding to higher values.

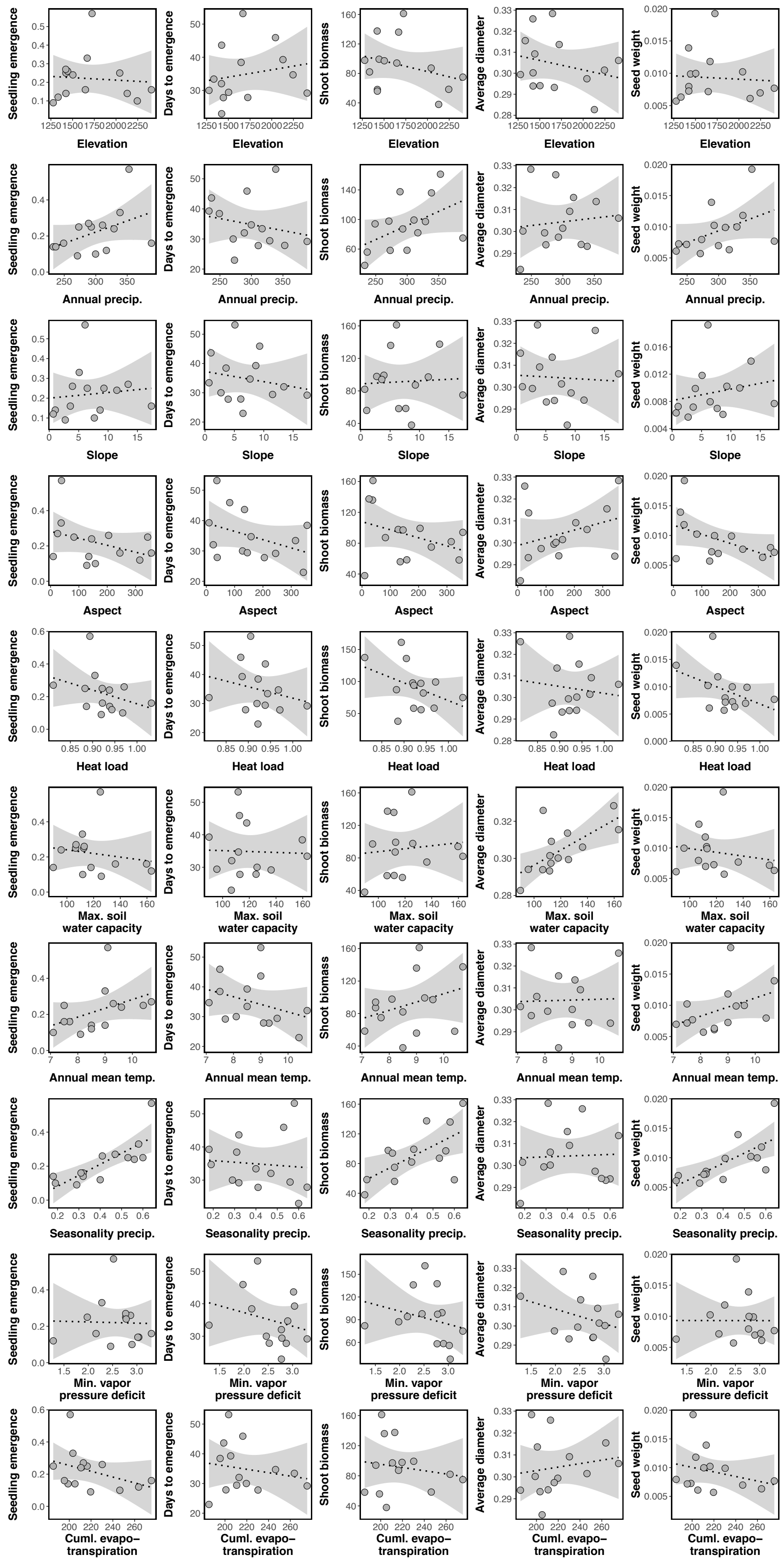


**Figure S2: Scatter plots depicting correlations for all pairs of environmental variables and mean phenotypic values across populations.**

| | AS | BM | BV | DC | DH | EW | FR | HO | JC | LV | MD | PL | PT | PW | SC | SS | VM |
|---|---|---|---|---|---|---|---|---|---|---|---|---|---|---|---|---|---|
| AS | 0 | 0.04 | 0.104 | 0.093 | 0.045 | 0.118 | 0.051 | 0.048 | 0.068 | 0.141 | 0.129 | 0.115 | 0.095 | 0.049 | 0.044 | 0.136 | 0.075 |
| BM | 0.012 | 0 | 0.081 | 0.07 | 0.034 | 0.093 | 0.055 | 0.04 | 0.049 | 0.113 | 0.104 | 0.089 | 0.072 | 0.054 | 0.048 | 0.11 | 0.053 |
| BV | 0.036 | 0.027 | 0 | 0.03 | 0.079 | 0.061 | 0.133 | 0.081 | 0.053 | 0.062 | 0.039 | 0.049 | 0.061 | 0.126 | 0.128 | 0.048 | 0.052 |
| DC | 0.032 | 0.022 | 0.009 | 0 | 0.068 | 0.062 | 0.119 | 0.07 | 0.047 | 0.068 | 0.045 | 0.053 | 0.059 | 0.114 | 0.113 | 0.054 | 0.047 |
| DH | 0.014 | 0.01 | 0.026 | 0.022 | 0 | 0.093 | 0.057 | 0.038 | 0.046 | 0.112 | 0.103 | 0.088 | 0.07 | 0.056 | 0.051 | 0.109 | 0.049 |
| EW | 0.043 | 0.031 | 0.02 | 0.02 | 0.032 | 0 | 0.148 | 0.103 | 0.072 | 0.062 | 0.067 | 0.052 | 0.068 | 0.143 | 0.142 | 0.06 | 0.063 |
| FR | 0.016 | 0.018 | 0.052 | 0.045 | 0.019 | 0.06 | 0 | 0.058 | 0.091 | 0.173 | 0.159 | 0.149 | 0.123 | 0.036 | 0.036 | 0.169 | 0.101 |
| HO | 0.015 | 0.012 | 0.027 | 0.023 | 0.012 | 0.036 | 0.019 | 0 | 0.051 | 0.121 | 0.107 | 0.097 | 0.08 | 0.056 | 0.052 | 0.115 | 0.058 |
| JC | 0.022 | 0.015 | 0.017 | 0.014 | 0.014 | 0.023 | 0.032 | 0.016 | 0 | 0.081 | 0.071 | 0.061 | 0.056 | 0.087 | 0.084 | 0.074 | 0.038 |
| LV | 0.054 | 0.041 | 0.02 | 0.022 | 0.041 | 0.02 | 0.075 | 0.045 | 0.027 | 0 | 0.061 | 0.041 | 0.067 | 0.167 | 0.167 | 0.049 | 0.068 |
| MD | 0.049 | 0.037 | 0.012 | 0.014 | 0.037 | 0.022 | 0.067 | 0.039 | 0.024 | 0.02 | 0 | 0.054 | 0.073 | 0.152 | 0.154 | 0.045 | 0.068 |
| PL | 0.041 | 0.03 | 0.015 | 0.016 | 0.03 | 0.016 | 0.06 | 0.034 | 0.019 | 0.012 | 0.017 | 0 | 0.05 | 0.141 | 0.143 | 0.045 | 0.048 |
| PT | 0.033 | 0.024 | 0.02 | 0.019 | 0.023 | 0.022 | 0.047 | 0.027 | 0.018 | 0.022 | 0.025 | 0.016 | 0 | 0.118 | 0.117 | 0.07 | 0.048 |
| PW | 0.015 | 0.018 | 0.049 | 0.042 | 0.018 | 0.057 | 0.011 | 0.018 | 0.031 | 0.071 | 0.063 | 0.056 | 0.045 | 0 | 0.038 | 0.162 | 0.097 |
| SC | 0.013 | 0.015 | 0.048 | 0.041 | 0.016 | 0.056 | 0.011 | 0.017 | 0.029 | 0.07 | 0.063 | 0.056 | 0.044 | 0.012 | 0 | 0.162 | 0.094 |
| SS | 0.052 | 0.039 | 0.015 | 0.017 | 0.039 | 0.019 | 0.072 | 0.042 | 0.025 | 0.015 | 0.014 | 0.014 | 0.023 | 0.068 | 0.067 | 0 | 0.066 |
| VM | 0.024 | 0.016 | 0.016 | 0.014 | 0.015 | 0.02 | 0.037 | 0.018 | 0.011 | 0.022 | 0.023 | 0.015 | 0.015 | 0.035 | 0.033 | 0.022 | 0 |

**Figure S3: Pairwise estimates of genetic differentiation across sampling localities.** Hudson's $F_{ST}$ is represented in the upper triangle and Nei's $D$ is represented in the lower triangle. Colors correspond to the magnitude of estimates, with warmer colors corresponding to higher values.

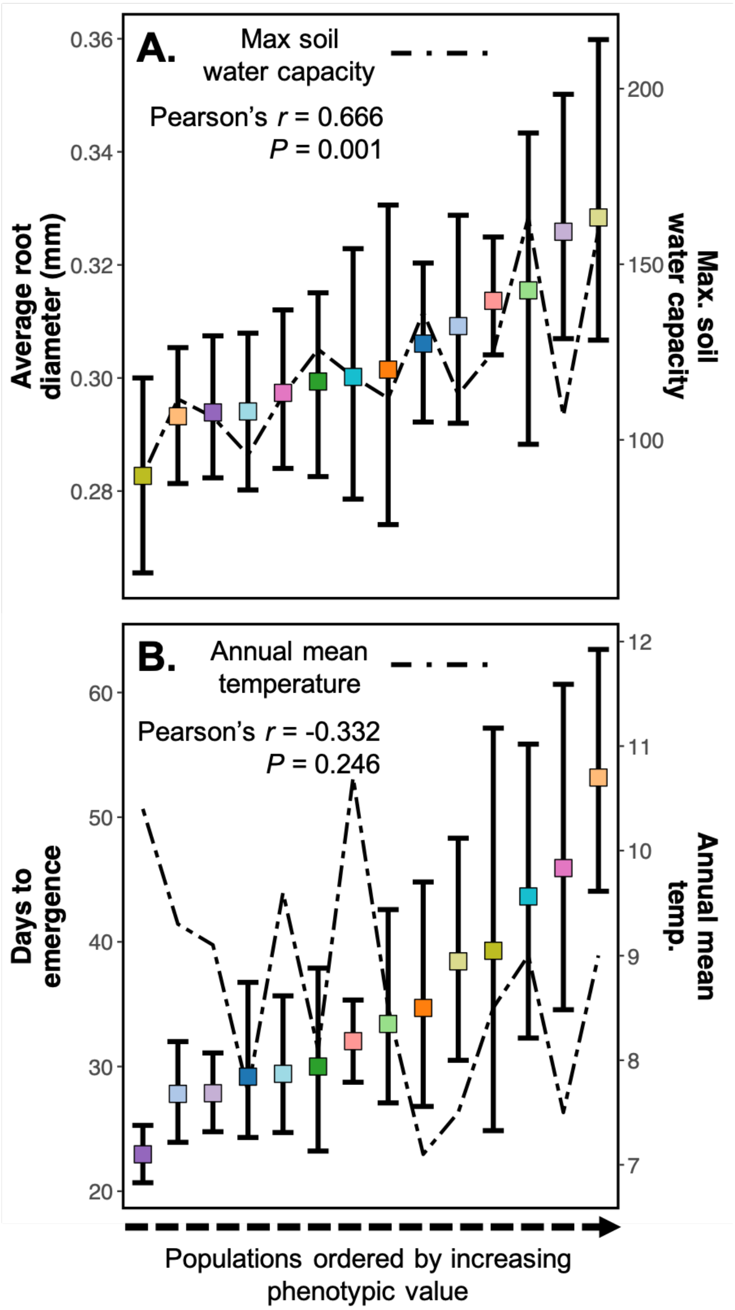


**Figure S4: Average root diameter and days to emergence data vary across populations and have mixed association (i.e., local adaptation) to environmental variables.** (**A**) and (**B**) Average root diameter and days to emergence vary by orders of magnitude among the populations. This suggests genetically-based differentiation of these two phenotypes across populations, although unmeasured maternal effects could also have contributed. Additionally, the association with strongest environmental correlation (described by Pearson's *r* and *P*-values) is represented by right y-axis and the line in each panel. The mean phenotype and 95% bootstrapped confidence intervals are represented by the squares and error bars. The populations are ordered by increasing mean phenotypic value. Colors are the same as those in the Fig. 1 map.

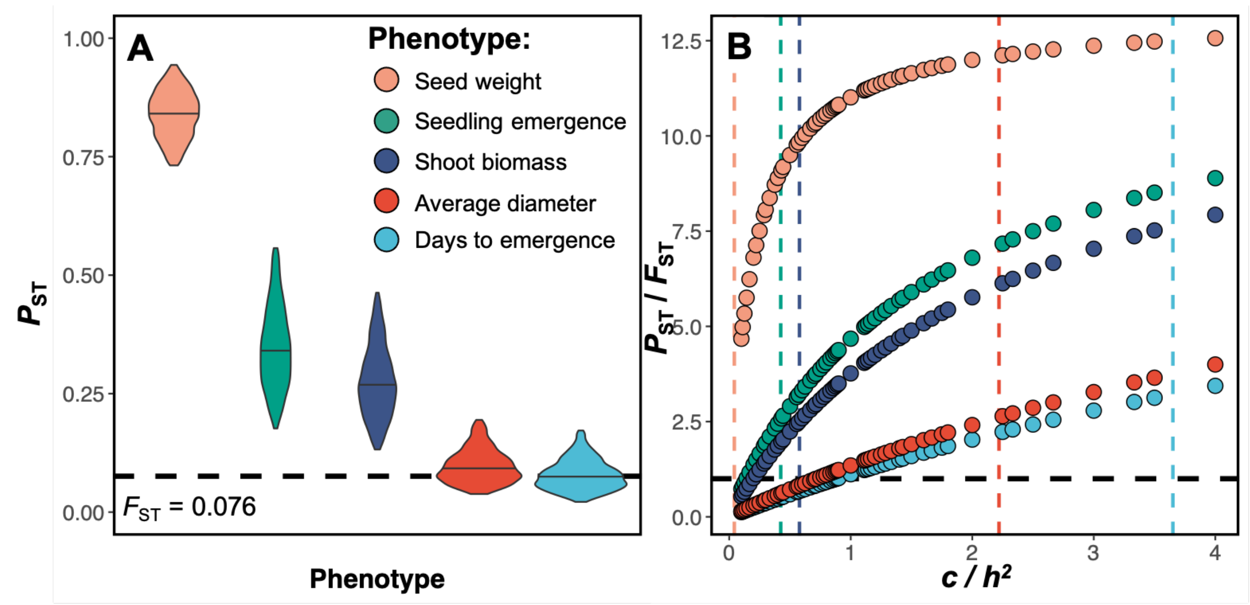


**Figure S5: $P_{ST}$ - $F_{ST}$ comparisons.** (**A**) $P_{ST}$ estimates calculated using $c/h^2 = 1$ for each phenotype. Violin plots represent the 95% credible interval from the posterior probability and the dashed line represents the multilocus $F_{ST}$ estimate. (**B**) $P_{ST}$ / $F_{ST}$ estimates for each phenotype at $c/h^2$ ratios ranging from 0.1–4.0. Horizontal dashed line represents where $P_{ST}$ and $F_{ST}$ are equal. Vertical dashed lines represent the critical value of the $c/h^2$ ratio (i.e., where the lower 95% CI of $P_{ST}$ equals the upper 95% CI of $F_{ST}$) for each phenotype. Phenotype is designated by color.

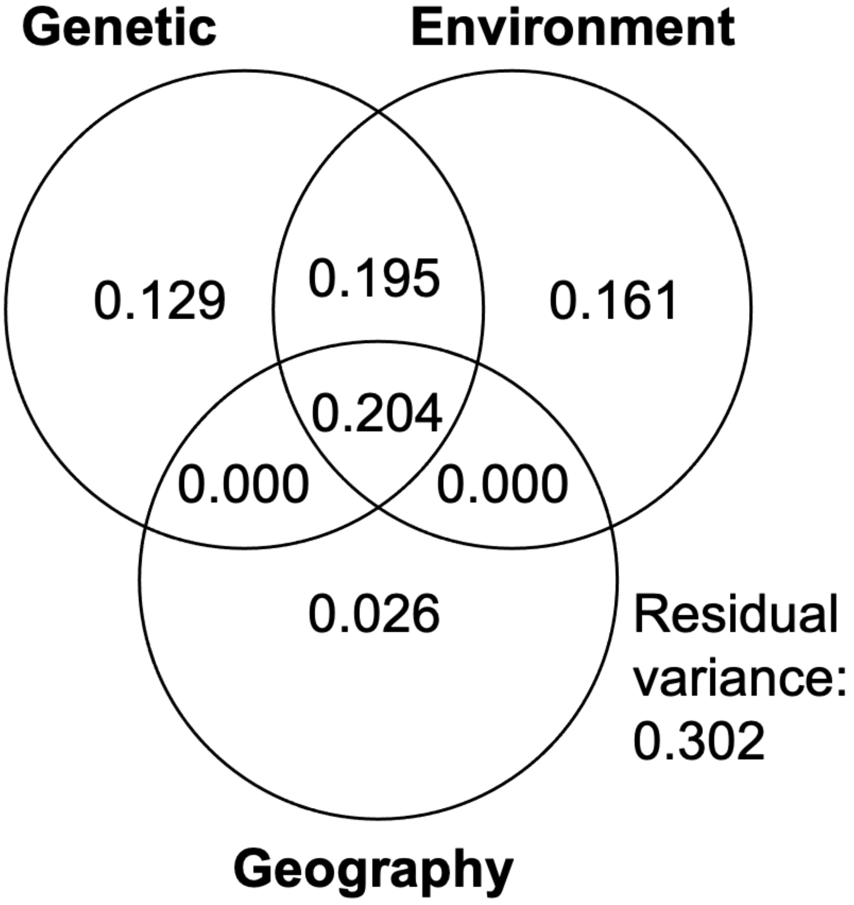


**Figure S6:** Venn diagram illustrating partial variance of genetic, environment, and geographic effects on phenotypic variation in terms of adjusted $r^2$. The total effect of all three predictors explained 69.84% of the total variance in phenotype in the common garden experiment.

### Supporting Information Tables

**Table S1: Phenotypic summaries for each population.** Provided as a csv file (10.5061/dryad.j0zpc86f4).

| *k* | chain 1 | chain 2 | chain 3 | chain 4 | mean |
|---|---|---|---|---|---|
| **2** | 35957053.4 | 34805863.1 | 34859146.2 | 34751716.8 | **35093444.9** |
| 3 | 60238272.3 | 58430430.0 | 55324174.3 | 62010775.1 | 59000913.0 |
| 4 | 69186354.8 | 69243142.9 | 74238907.7 | 74523224.4 | 71797907.5 |
| 5 | 88135378.4 | 81264483.1 | 80201813.9 | 78505312.0 | 82026746.8 |
| 6 | 88003037.4 | 86475339.2 | 99617573.1 | 92627979.5 | 91680982.3 |
| 7 | 125840808.6 | 116920867.1 | 204622515.3 | 114634552.3 | 140504685.8 |
| 8 | 205579857.1 | 217789135.9 | 172167727.9 | 1861692605.7 | 614307331.6 |

**Table S2: Deviance information criterion (DIC) values for different ancestral clusters (*k*) outputted from ENTROPY.** *k* = 2 model fit the data best indicated by lowest DIC value and presented in bold.

**Table S3: Environmental variable summaries for each population.** Provided as a csv file (10.5061/dryad.j0zpc86f4).

| Phenotype | $c/h^2$ | $P_{ST}$ | Lower 95% CI | Upper 95% CI | Critical $c/h^2$ | $P_{ST}$ - $F_{ST}$ |
|---|---|---|---|---|---|---|
| Seedling emergence | 1 | 0.354 | 0.557 | 0.177 | 0.424 | $P_{ST} > F_{ST}$ |
| Seed weight | 1 | 0.834 | 0.944 | 0.732 | 0.042 | $P_{ST} > F_{ST}$ |
| Days to emergence | 1 | 0.085 | 0.172 | 0.021 | 3.651 | $P_{ST} = F_{ST}$ |
| Shoot biomass | 1 | 0.285 | 0.463 | 0.132 | 0.578 | $P_{ST} > F_{ST}$ |
| Average diameter | 1 | 0.102 | 0.195 | 0.038 | 2.220 | $P_{ST} = F_{ST}$ |

**Table S4: $P_{ST}$ - $F_{ST}$ comparison for each phenotype.** Reported values represent the mean and 95% credible interval of $P_{ST}$ calculated when $c/h^2 = 1$. $P_{ST}$ - $F_{ST}$ comparison was decided if the 95% credible interval was wholly greater than, less than, or encompassed the multilocus $F_{ST}$ of 0.076.
